# Pan-cancer characterization and clinical outcome of ecDNA-enhancer mediated transcriptional dysregulation

**DOI:** 10.1101/2023.09.06.556610

**Authors:** Tengwei Zhong, Huan zhao, Yupeng Liu, Danyang Yan, Yuguo Li, Zhiyun Guo

## Abstract

Extrachromosomal circular DNAs (ecDNAs) are extrachromosomal circular DNA elements harboring enhancers that can function as mobile transcriptional enhancers to promote oncogene amplification in diverse human cancers. However, little is known about the mechanism how ecDNA and relevant enhancers influence the transcription control in different cancers. Here, we performed a comprehensive pan-cancer analysis to explore the potential transcriptional regulatory network between ecDNA and enhancer, and the association with clinical outcomes in different human cancers. Firstly, we identified 18,385 enhancers on ecDNA in 1,921 samples of 20 TCGA tumors. By integrating corresponding RNA-seq data, we found that the expression levels of enhancer RNAs (eRNAs) were significantly different between ecDNA regions and corresponding linear chromosomes. This suggested that enhancers on ecDNA may employ eRNA to perform transcriptional regulation. Furthermore, we identified enhancer co-amplification with oncogenes and found that enhancers on ecDNA significantly affect the expression of co-amplified cancer genes and consequently influence the survival of cancer patients. In addition, we found that, compared to corresponding linear chromosomes, enhancers on ecDNA generally have lower levels of methylation, which may be mainly caused by the enhancer co-amplification with oncogenes. Genome-wide analysis of open chromatin regions revealed significantly higher levels of chromatin accessibility and certain chromosomal preferences in ecDNA regions comparing corresponding linear chromosomes. Moreover, we found that the downregulation of antigen presentation genes and the suppression of antigen presentation pathways may cause the tumor immune evasion mediated by ecDNA. Finally, we identified tumor-infiltrating activated mast cells in ecDNA-positive patients, which are associated with poorer prognosis. In summary, through pan-cancer analysis, we illustrate the characteristics and regulatory mechanisms of ecDNA and its enhancers in cancers, providing insights for future cancer treatment.

## 1. Introduction

Extrachromosomal DNA (ecDNA) is a circular DNA molecule that is located outside of the chromosomes, which has been discovered in around 40% of cancers [1]. Due to the absence of centromeres, ecDNA can undergo uneven segregation during cell division, leading to the high heterogeneity of cancer cells. According to recent studies, the high copy number amplification of ecDNA leads to significantly increased transcriptional expression levels of oncogenes that are highly enriched on ecDNA, which may be a primary factor driving the occurrence and development of tumors. [2].

Enhancers are DNA elements that are located near the promoter regions of target genes and regulate the expression of target genes in a spatial and temporal manner [3]. Studies have shown that most oncogenes are controlled by enhancers capable to transcribe enhancer RNA (eRNA) or enhancer clusters located in locus control regions [4], and they further promote the initiation and progression of tumors. In 2019, Andrew R. Morton et al. demonstrated that EGFR preferentially co-amplifies with two active endogenous enhancer elements, and enhancer co-amplification was observed in multiple tumors. The endogenous enhancers on the amplicon promote the proliferation of cancer cells [5]. Subsequent studies have also revealed that ectopic enhancer hijacking can compensate for the loss of local gene regulatory elements [6]. Further research has found that ecDNA can act as a mobile super-enhancer, driving amplification of genome-wide transcription [7], and ecDNAs can “cluster” together becoming ecDNA hubs, which can drive cooperative oncogene expression by strengthening intermolecular enhancer signals [8]. Increasing reports have emerged on the involvement of these enhancers in ecDNA regulation and their impact on cancer initiation and progression [9–12]. Although the importance of enhancers on ecDNA has been reported previously, it remains unclear whether enhancers on ecDNA have common or specific features in cancers, and whether the enhancer RNA on ecDNA plays a significant role in tumor regulation. Therefore, this study comprehensively analyzed the characteristics and functions of ecDNAs, the relevant regulatory elements or enhancers, and eRNAs in various tumors, from a pan-cancer perspective by integrating multi-omics data.

We integrated ecDNA data and enhancers from patients with various tumors in TCGA and identified enhancers on ecDNA in multiple tumors. By integrating enhancer expression data, we found that eRNA expression levels were significantly higher in ecDNA regions compared to corresponding linear chromosomes. We identified a large number of enhancer-oncogene co-amplification segments in multiple tumors and found that frequently co-amplified oncogenes such as EGFR and SOX2 were more highly expressed in ecDNA-positive (ecDNA+) patients who also had worse prognosis. In addition, methylation analysis revealed significant differences in methylation levels between ecDNA regions and corresponding linear chromosomes, with enhancers on ecDNA exhibiting lower methylation. By integrating open-access data from TCGA, we found that ecDNA regions had higher chromatin accessibility and recruited transcription factors involved in tumor biological processes. This indicated that the transcription factors in the open chromatin regions of ecDNA were mostly associated with transcriptional regulation, such as DNA-binding transcription activator activity. Finally, tumor immune analysis revealed immune evasion associated with ecDNA in pan-cancer. Decreased expression of antigen presentation genes and downregulation of the antigen presentation pathway were the main factors contributing to immune evasion. In conclusion, through pan-cancer analysis, we elucidated the characteristics and regulatory mechanisms of ecDNA and its enhancers in cancer, providing insights for future tumor-related therapies.

## 2. Materials and Methods

### 2.1. Identification of ecDNA Enhancers and Pan-cancer Analysis of eRNA Expression

The ecDNA (hg19) regions of TCGA tumor samples predicted by the AA algorithm were downloaded from a previous study by Hoon Kim et al. [13]. The enhancer information and expression data for corresponding TCGA samples were downloaded from the study by Han Chen et al. [14]. Enhancers that completely overlapped with the ecDNA regions were defined as enhancers on ecDNA using the intersect function in bedtools v2.29.2. Enhancers with no expression in the ecDNA region (RPKM>0) were filtered out. For the comparison between ecDNA+ and ecDNA- samples, the average expression levels of enhancers in both samples were calculated respectively. For the comparison between ecDNA regions and corresponding linear chromosome regions, the expression levels of enhancers in both regions were analyzed respectively.

### 2.2. Identification of ecDNA Enhancer-Oncogene Co-amplification

The oncogene information was obtained from the ONGene database (http://www.ongene.bioinfo-minzhao.org/). TCGA tumor gene expression data and copy number segment data were obtained from UCSC Xena (https://xena.ucsc.edu/). The intersect function in bedtools v2.29.2 was used to identify the co-amplification of enhancers and cancer genes that are completely located on the same ecDNA fragment. Afterwards, a filtering process is applied to exclude co-amplified fragments with copy number less than 2. The risk factor association analysis for these oncogenes was calculated using the R packages “survival” and “ggrisk”.

### 2.3. Methylation Analysis of ecDNA

The methylation data of TCGA tumor samples were downloaded from UCSC Xena (https://xena.ucsc.edu/). The intersect function in bedtools v2.29.2 was used to capture the methyl ation sites in the ecDNA regions or enhancer regions on ecDNA.

### 2.4. Chromatin Accessibility Analysis

The ATAC data was obtained from the study by Howard Y. Chang et al [15], and the LiftOver tool was used to convert the ATAC regions from hg38 to hg19. The intersect function in bedtools v2.29.2 was used to capture the ATAC peaks in the ecDNA regions or enhancer regions on ecDNA. For the analysis of accessibility features, the coverage function (default parameters -a and -b) in bedtools v2.29.2 was used to calculate the ATAC-seq peak density of 22 pairs of autosomal chromosomes and one pair of sex chromosomes, and ecDNA regions in each tumor. When the ATAC-seq peak density of the complete chromosome regions was lower than that of the corresponding ecDNA regions, it was defined as “High ATAC density of ecDNA”. In contrast, it was defined as “Low ATAC density of ecDNA”.

### 2.5. Identification of Transcription Factor on ecDNA and Motif Analysis

The getfasta function (default parameters -fi and -bed) in bedtools v2.29.2 was used to obtain sequence of all accessible regions in the ecDNA. The MEME v5.5.2 [16] streme algorithm was employed to identify motifs. The identified motifs were then compared to the transcription factor database JASPAR (https://jaspar.genereg.net/) using the tomtom tool. Motifs with an E value < 0.05 and p.adjust < 0.05 were considered as threshold for selecting transcription factors binding in the accessible regions of the ecDNA. Gene Ontology (GO) enrichment was analyzed using the R package “clusterProfiler”.

### 2.6. Analysis of ecDNA Tumor Immune Infiltration

The CIBERSORT algorithm was utilized to calculate the proportion of 22 immune cells in each patient (with the parameter perm = 1000). Samples with a significance level of p < 0.05 were selected to obtain immune infiltration data. The difference in immune cell infiltration between ecDNA+ and ecDNA- samples was evaluated using the paired Wilcoxon test.

### 2.7. Immunological Checkpoint and Antigen Presentation Analysis

Thirteen common immunological checkpoints were obtained from previous studies [17–18]. The antigen presentation pathways (REACTOME_CLASS_I_MHC_MEDIATED_ANTIGEN_ PROCESSING_PRESENTATION, REACTOME_MHC_CLASS_II_ANTIGEN_PRESENTATI ON, GOBP_ANTIGEN_PROCESSING_AND_PRESENTATION_OF_PEPTIDE_ANTIGEN_V IA_MHC_CLASS_I, GOBP_ANTIGEN_PROCESSING_AND_PRESENTATION_OF_PEPTIDE_OR_POLYSACCHARIDE_ANTIGEN_VIA_MHC_CLASS_II) and their corresponding anti gen presentation genes were obtained from the MSigDB (https://www.gsea-msigdb.org/gsea/msigdb/index.jsp) database. The expression matrices of immune checkpoint genes and antigen presentation genes were extracted from the previously downloaded gene expression data using R v3.6.1. And the expression levels of immune checkpoint genes and antigen presentation genes in ecDNA+ and ecDNA- were analyzed respectively. The GSVA score of the antigen presentation pathway was calculated using the R package “GSVA”.

### 2.8. Construction of Immune Cell Signature

The clinical data of TCGA patients were obtained using the R package “TCGAbiolinks”. Immune cells related to the patient lifecycle were initially screened using univariate COX regression analysis (P<0.05). The LASSO model was then employed to select prognostic signatures associated with patient survival using a punishment mechanism.

### 2.9. Survival Analysis

All survival analysis was calculated by R package “survival”.

### 2.10. Statistical analysis

Statistical analyses were performed using R v3.6.1 statistical software. The Wilcoxon test a nd Wilcoxon paired test were performed to make group comparison. Bedtools shuffle was utilized for random sampling, generating a total of 1000 groups. The sampled regions hav e the same size as ecDNA regions and do not overlap with them. All correlation analyses were performed using the Spearman method. We used the following convention for symbols indicating statistical significance: *: P≤0.05, **: P≤0.01, ***: P≤0.001, ****: P≤0.0001, ns: P>0.05.

## 3. Results and Discussion

### 3.1. Identification of ecDNA Enhancers in TCGA Tumors

Previous studies have suggested that ecDNA harboring enhancers can serve as a mobile regulatory element, which regulates the expression of oncogenes in the form of individual ecDNAs or ecDNA hubs. To investigate the characteristics of ecDNA enhancers in tumors from a pan-cancer perspective, we conducted the following work: First, we downloaded 3,014 ecDNAs from 1,921 samples of 20 TCGA tumors reported in the study by Hoon Kim et al [2]. To better distinguish patient samples, we defined patients with ecDNA as “ecDNA-positive” (ecDNA+), and patients without ecDNA as “ecDNA-negative” (ecDNA-). Statistics on these 20 tumor patients revealed that the proportion of ecDNA+ samples was lower than ecDNA- samples in the entire sample set, and most ecDNA+ samples contained fewer than 10 ecDNAs each (Supplementary Figure S1). Combined with clinical data, we found that the average age of the majority of ecDNA+ patients was higher than that of ecDNA- patients (Figure 1B), and there was a higher proportion of male patients than female patients (Supplementary Figure S2). Furthermore, ecDNA+ patients had a significantly higher proportion in the advanced stages (particularly stage IV) compared to ecDNA- patients (Figure 1C). Taken together, it is indicated that ecDNA tends to be present in tumors with a higher malignant grade.

**Figure 1.**
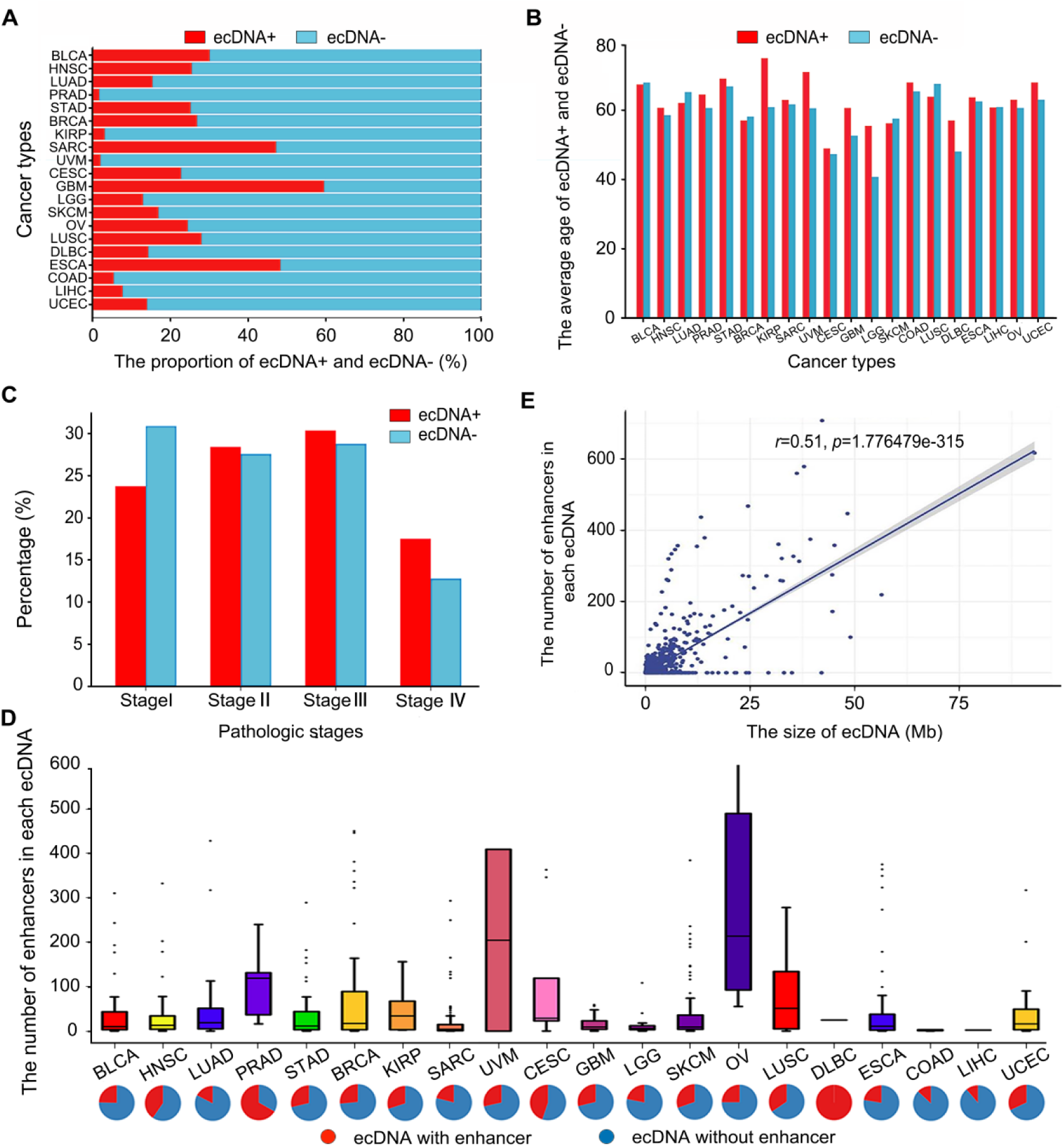
Statistical analysis of ecDNAs and enhancers on ecDNA in 20 tumors. A. Proportion of ecDNA+ and ecDNA- patients among the total patients. B. Distribution of average age among ecDNA+ and ecDNA- patients. C. Proportion of pathological stages in ecDNA+ and ecDNA- patients. D. The bar graph indicates the number of enhancers on the ecDNA regions; and the pie graph indicates the proportion of ecDNA harboring enhancers compared to the total number of ecDNA. E. Spearman correlation between ecDNA size and the number of enhancers.

In addition, to identify the enhancers on ecDNA in these 20 tumors, we integrated the enhancers identified by Han Chen et al [14] in these 20 TCGA tumors, resulting in a total of 18,385 enhancers on ecDNA. The statistical analysis showed that ovarian serous cystadenocarcinoma (OV) had the highest average number of enhancers on ecDNA (an average of 294 enhancers per ecDNA), followed by uveal melanoma (UVM) (an average of 205 enhancers per ecDNA) (Figure 1D). In terms of the proportion of ecDNAs containing enhancers in each tumor, prostate adenocarcinoma (PRAD) had 70% of ecDNAs harboring enhancers, followed by cervical squamous cell carcinoma (CESC) with 45%. Moreover, Spearman correlation analysis revealed a significant positive correlation between the number of enhancers and the number of ecDNAs in the tumor (p=2.8e-2, r=0.24, Supplementary Figure S3), as well as the size of ecDNA (p=1.776479e-315, r=0.51, Figure 1E).

### 3.2. Significantly Different Expression Levels of eRNAs in ecDNA Regions Compared to the Corresponding Linear Chromosomes

Previous studies have shown that some enhancers can transcribe eRNAs, which significantly affect their activity [14]. To investigate the eRNAs associated with enhancers on ecDNA, we identified the eRNA in 20 tumors. Due to the fact that the expression of most enhancers (over 95%) in UCEC was zero and LIHC had a very low number of enhancers on ecDNA, we excluded these two tumor types from all subsequent analyses. Finally, we obtained 6,959 eRNAs on ecDNA in 18 tumors (Supplementary Table 1). We performed a paired rank sum test to compare eRNA expression differences between ecDNA+ and ecDNA- samples in the 18 tumors (Figure 2A). It is indicated that, except for BRCA, there were significant differences in eRNA expression between ecDNA+ and ecDNA- samples in 17 tumors: the expressions of eRNAs were upregulated in 15 ecDNA+ samples and downregulated in 2 ecDNA+ samples when compared to ecDNA- samples. It is suggested that eRNAs play an important role in enhancer-mediated ecDNA regulation in tumors.

**Figure 2.**
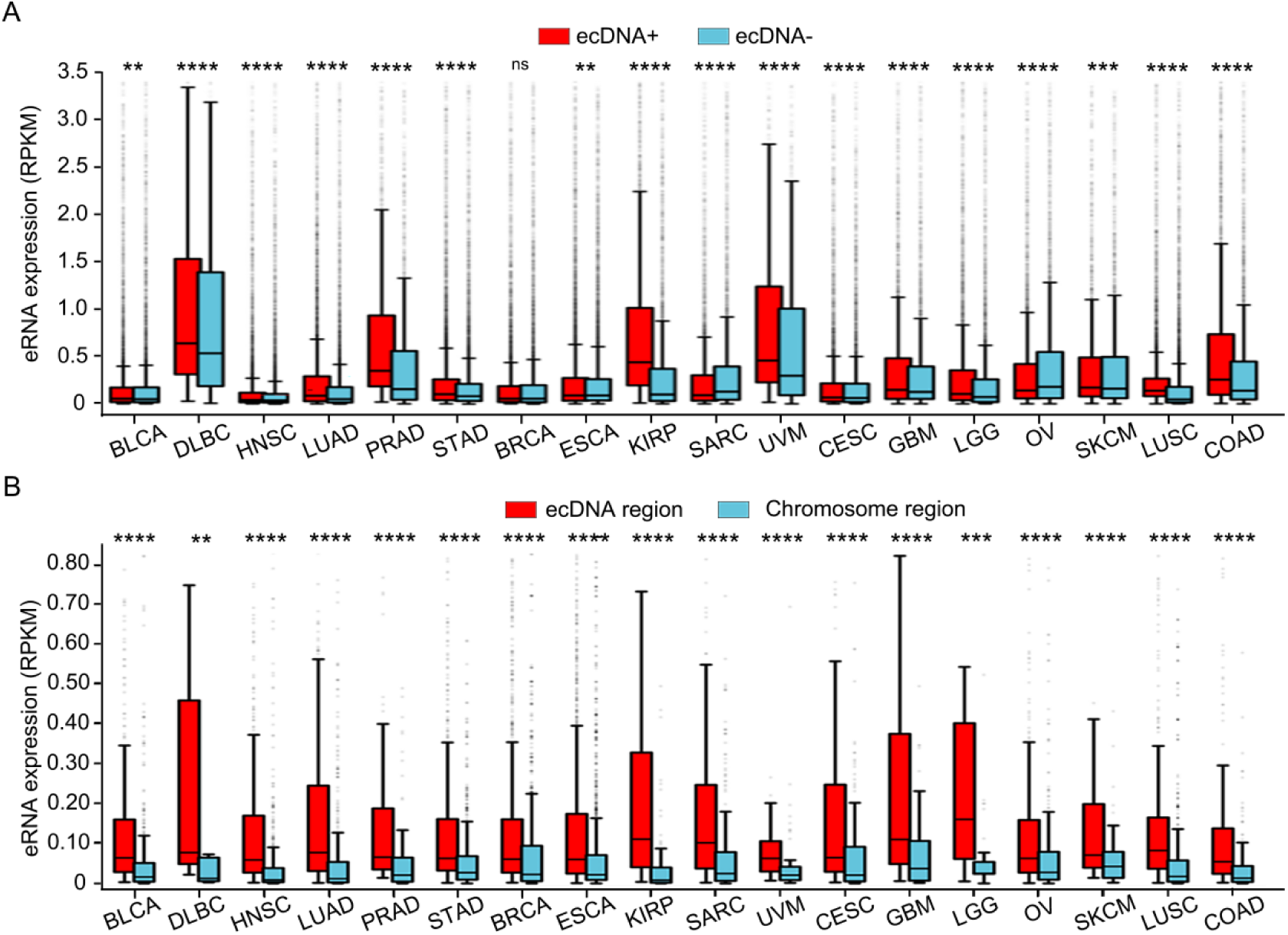
Analysis of eRNA expression in ecDNA+/ecDNA- samples and ecDNA regions in 18 tumors. A. Expression of enhancers in ecDNA+ and ecDNA- patients. B. Expression of enhancers in the ecDNA regions and the corresponding linear regions. The significance was determined using a Wilcoxon test (*p≤0.05, **p≤0.01, ***p≤0.001, nsp>0.05).

To determine whether the differences in eRNA expression between ecDNA+ and ecDNA- samples were caused by the ecDNA regions, we analyzed the eRNA expression in the ecDNA regions of ecDNA+ samples compared to the corresponding linear regions of ecDNA- samples in 18 tumors. Notably, in these tumors, the eRNA expression in the ecDNA regions was significantly higher than in the corresponding linear regions. The most significant higher expression of the eRNA was found in the ESCA (p=2.08E-117) (Figure 2B). Previous studies have reported that specific active enhancers are essential for the survival of ESCA cancer cells [19].

Furthermore, to investigate whether the eRNA expression differences were specifically caused by the ecDNA regions, we examined the expression levels of eRNAs in the ecDNA regions and the regions outside of ecDNA on the same chromosome, as well as the corresponding linear regions. We performed 1,000 random sampling analyses on these regions (Supplementary Figure S4). The results showed that the expression levels of enhancers in the ecDNA regions were significantly higher than in the regions outside of ecDNA, whereas there was no significant difference in enhancer expression between the regions outside of ecDNA and the most corresponding linear regions. Taken together, these results indicated significant differences in eRNA expression between ecDNA+ and ecDNA- samples, primarily driven by eRNAs in the ecDNA regions. It is suggested that ecDNA enhancers may play an important role in tumor regulation through eRNA expression.

### 3.3. Analysis of Enhancer-Oncogene Co-Amplification in ecDNA

Co-amplification of enhancers and oncogenes on ecDNA can lead to high expression of oncogenes, thereby affecting cancer fitness [20]. To analyze the characteristics of enhancer-oncogene co-amplification from a pan-cancer perspective, we utilized the cancer copy number data from UCSC Xena to identify enhancer-oncogene co-amplifications in 18 tumors. Finally, we identified a total of 15,134 enhancer-oncogene co-amplifications in 16 tumors (Supplementary Table 2). Among them, BRCA had the highest number of oncogene-enhancer co-amplifications (4,676 pairs), followed by ESCA with 2,045 pairs. The frequent utilization of oncogenes in co-amplification often implies more crucial functions. Moreover, we assessed the frequently utilized oncogenes in co-amplifications (Figure 3A). As expected, oncogenes such as EGFR, SOX2, TRIB1, MYC, and PVT1, which have been widely reported to be involved in ecDNA-mediated tumor regulation, were frequently utilized in co-amplifications [2, 8].

**Figure 3.**
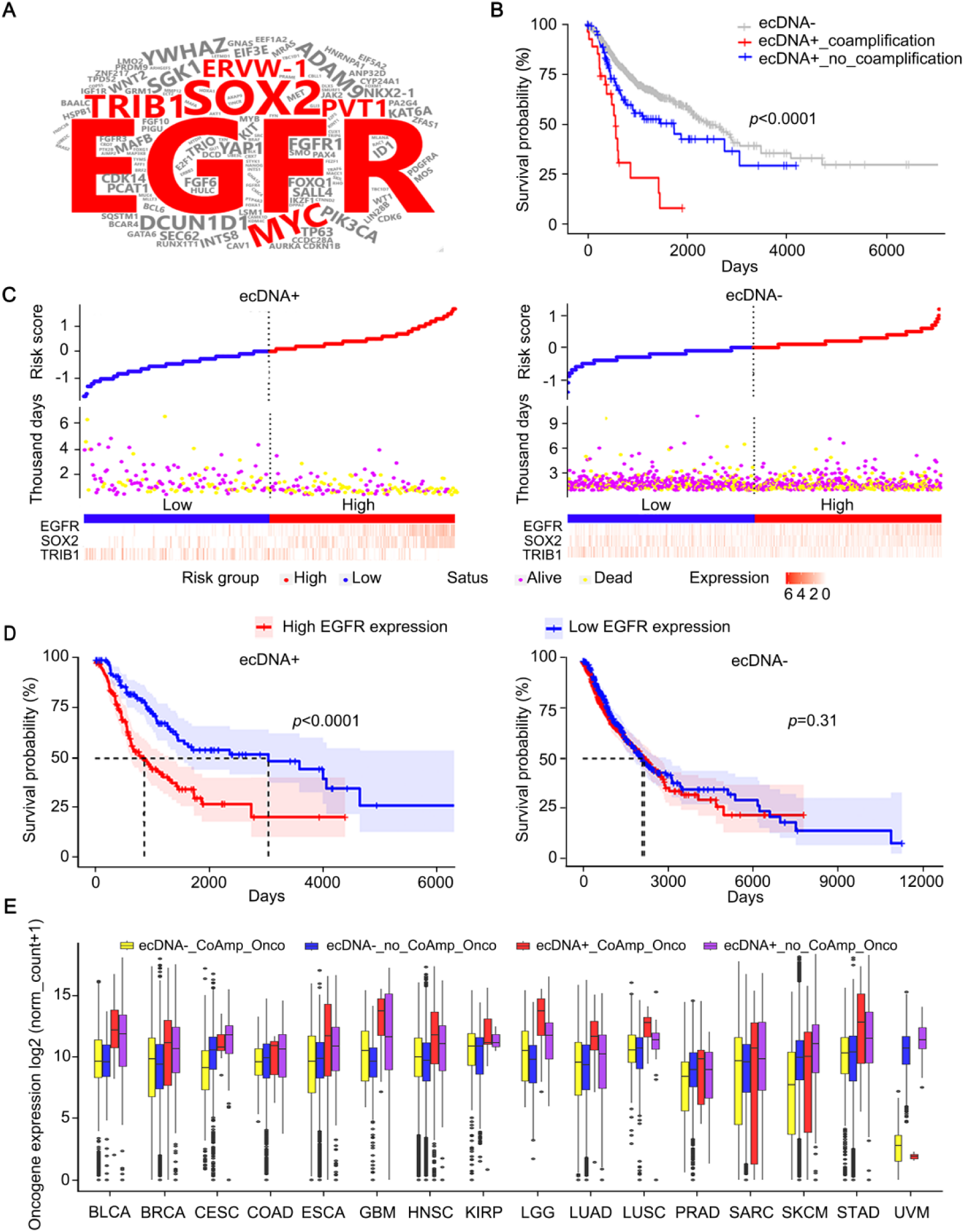
Analysis of enhancer-oncogene co-amplification in ecDNA. A. Frequently co-amplified oncogenes. The size of the oncogene names represents the frequency of their involvement in co-amplification. B. Impact of EGFR on the prognosis of ecDNA-, ecDNA+ with co-amplification, and ecDNA+ without co-amplification patients. C. Risk factor associations in ecDNA+ and ecDNA-patients. Dashed line represents the median risk score, blue and red blocks represent low-risk and high-risk patients, respectively. Yellow dots represent deceased patients, and purple dots represent survived patients. D. Effect of EGFR expression levels on the prognosis of ecDNA+ and ecDNA- patients. E. Expression of oncogenes involved in co-amplification and oncogenes not involved in co-amplification in ecDNA+ and ecDNA- samples. ecDNA+_CoAmp_Onco: Oncogenes involved in co-amplification in ecDNA+; ecDNA+_no_CoAmp_Onco: Oncogenes not involved in co-amplification in ecDNA+; ecDNA-_CoAmp_Onco: Oncogenes involved in co-amplification in ecDNA-; ecDNA-_no_CoAmp_Onco: Oncogenes not involved in co-amplification in ecDNA-.

To investigate whether these enhancer-oncogene co-amplifications have an impact on patient outcomes, we analyzed the survival of ecDNA+ patients with co-amplified oncogenes compared to ecDNA+ patients without co-amplifications. The results revealed that ecDNA+ patients with co-amplified oncogenes had significantly worse survival compared to ecDNA+ patients without co-amplifications (P<0.0001), and both groups had lower survival rates than ecDNA- patients (Figure 3B and Supplementary Figure S5), suggesting that enhancer-oncogene co-amplifications affect patient outcomes.

Furthermore, we examined the expression and risk associations of the top three frequently involved co-amplified oncogenes (EGFR, SOX2, TRIB1) in ecDNA+ and ecDNA- patients, as well as their impact on patient survival. The results indicated that low-risk patients among the ecDNA+ group had longer survival compared to high-risk patients. EGFR, SOX2, and TRIB1 oncogenes exhibited significant expression spectrum characterization in high- and low-risk groups, particularly with high expression of EGFR and SOX2 in high-risk patients, which was not observed in ecDNA- patients (Figure 3C). We then performed survival analysis for these three genes, and the results indicated that the expression levels of these genes significantly influenced patient survival compared to ecDNA- patients (Figure 3D and Supplementary Figure S6).

In addition, to explore whether enhancers significantly affect the expression of co-amplified oncogenes, we classified oncogenes into four categories: 1) ecDNA+ co-amplified oncogenes (ecDNA+ CoAmp_oncogene); 2) ecDNA+ oncogenes without co-amplification (ecDNA+ no CoAmp_oncogene); 3) ecDNA- co-amplified oncogenes (ecDNA- CoAmp_oncogene); and 4) ecDNA- oncogenes without co-amplification (ecDNA- no CoAmp_oncogene). We then compared the expression levels of these four types of oncogenes. The results showed that in 15 out of 16 tumors (except UVM), the expression levels of ecDNA+ CoAmp_oncogenes were significantly higher than those of ecDNA- CoAmp_oncogenes (Figure 3E), with the highest significance found in LGG (p<2e-16). And the expression levels of ecDNA+ no CoAmp_oncogenes were significantly higher than those of ecDNA- no CoAmp_oncogenes in all 16 tumors, with the highest significance found in UVM (p<2e-16). We also found that in ecDNA+ samples, the expression levels of ecDNA+ CoAmp_oncogenes mostly higher than those of ecDNA+ no CoAmp_oncogenes, indicating the importance of co-amplification in influencing oncogene expression in tumors. Notably, we found a significant positive correlation between the frequency of oncogenes involved in co-amplification and the difference in their expression levels between ecDNA+ CoAmp_oncogene and ecDNA- CoAmp_oncogene (Supplementary Figure S7; Supplementary Table 3). Taken together, enhancer-oncogene co-amplifications in ecDNA+ patients significantly impact their survival, and enhancers involved in co-amplification significantly upregulate the expression of oncogenes.

### 3.4. Lower Methylation Commonly Found in the ecDNA Regions, Most Occurring in ecDNA+ Samples with Co-Amplification

Methylation plays a crucial regulatory role in tumor function and development. We examined the methylation levels of ecDNA+ and ecDNA- samples in 17 out of 18 tumors (except KIRP due to lack of TCGA methylation data). The results showed significant methylation differences in all 17 tumors, indicating that the presence or absence of ecDNA can impact the overall genomic methylation levels. Among these 17 tumors, methylation levels were significantly downregulated in ecDNA+ samples compared to ecDNA- samples in 10 tumors, while they were upregulated in 7 tumors (Supplementary Figure S8). This suggests that the presence of ecDNA can significantly impact the overall genomic methylation level in patients, but not all tumors exhibit a decrease in methylation levels in the presence of ecDNA.

We then conducted the analysis of methylation levels in the ecDNA regions of ecDNA+ samples and the corresponding linear regions of ecDNA- samples in the 17 tumors. The results showed that, except for HNSC and OV, the ecDNA regions exhibited lower levels of methylation compared to the corresponding linear regions in the remaining 15 tumors (Figure 4A). The results from random testing indicated that the overall methylation level in ecDNA regions was significantly lower compared to other random regions outside of ecDNA. In contrast, there was mostly no significant difference in methylation levels between the regions outside of ecDNA and the corresponding linear regions (Supplementary Figure S9). It is suggested that the lower methylation found in the ecDNA regions compared to the corresponding linear regions is not a random occurrence

**Figure 4.**
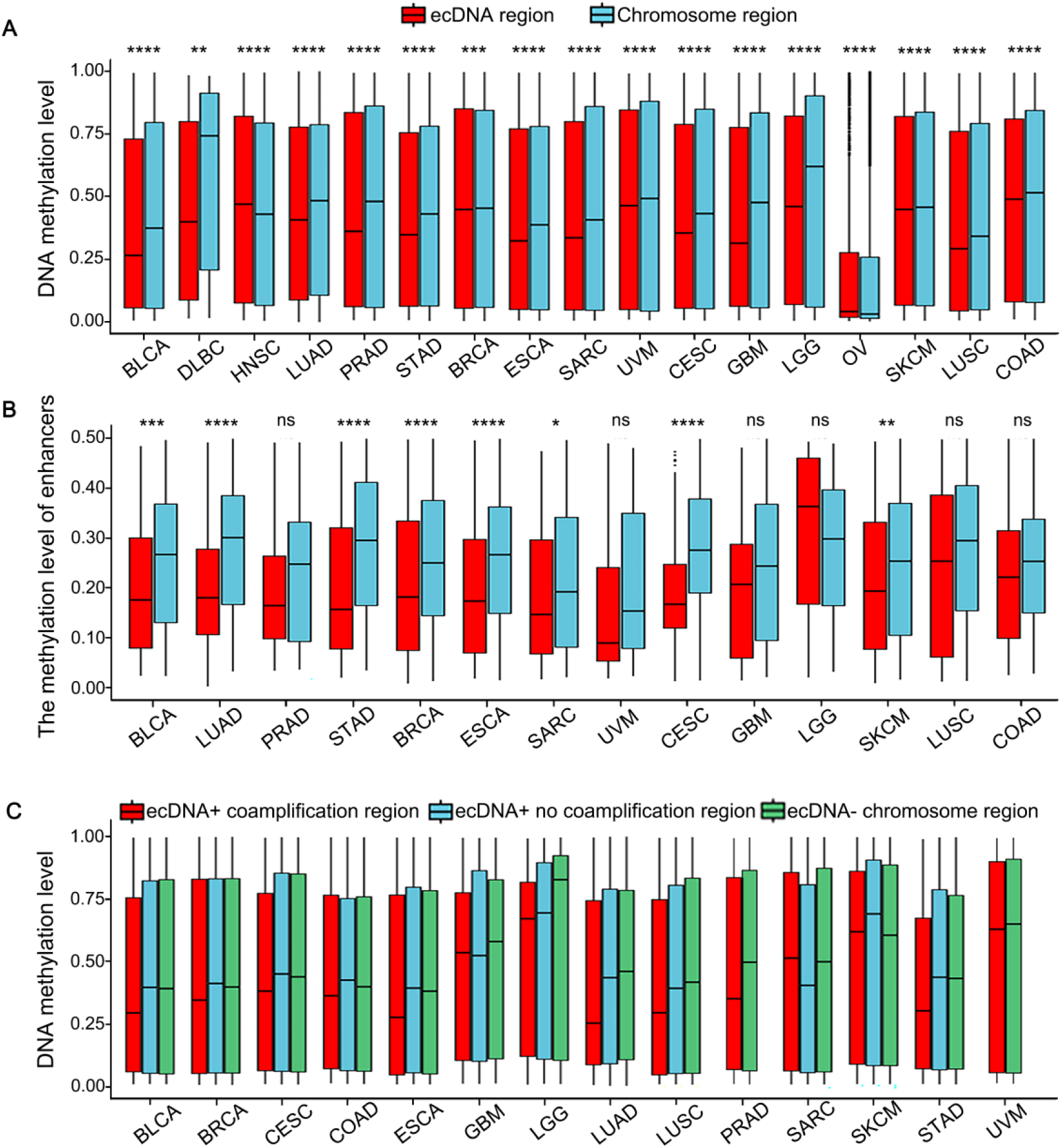
Methylation levels of ecDNA+ and ecDNA- samples, as well as ecDNA regions and the corresponding linear chromosome regions. A. Methylation levels of ecDNA regions and the corresponding linear regions in 17 tumors. The significance was determined using the paired Wilcoxon test (*p≤0.05, **p≤0.01, ***p≤0.001, nsp>0.05). B. Methylation levels of enhancers on ecDNA regions and the corresponding linear regions in 14 tumors. The significance was determined using the Wilcoxon test (*p≤0.05, **p≤0.01, ***p≤0.001, nsp>0.05). C. Methylation levels in the corresponding linear chromosome regions of ecDNA, ecDNA+ coamplification regions, and ecDNA+ no coamplification regions in 14 tumors. ecDNA+ coamplification regions refer to the co-amplified fragments in ecDNA+ samples; ecDNA+ no coamplification regions refer to the non-co-amplified fragments in ecDNA+ samples; ecDNA- chromosome regions refer to the corresponding linear chromosome regions in ecDNA- samples.

Previous studies have shown that one of the causes for enhanced transcription of oncogenes is the lower DNA methylation levels in the regulatory elements on ecDNA compared to chromosomal DNA [21]. Therefore, we analyzed the methylation levels of enhancers in the ecDNA regions and the corresponding linear regions in different tumors. Due to the lack of corresponding enhancer methylation data in DLBC, we analyzed 14 tumors out of these 15 tumors. The results showed that the methylation levels of enhancers in the ecDNA regions were lower than those in the corresponding linear regions in all 14 tumors. Among them, 8 tumors showed significant differences, with STAD and ESCA showing the highest significance, with p-values of 6.6e-07 and 2.2e-07, respectively (Figure 4B). Taken together, most tumors exhibit lower levels of methylation in ecDNA regions and enhancer regions on ecDNA compared to the corresponding linear regions.

Furthermore, we aimed to investigate whether enhancer-oncogene co-amplification is a factor contributing to the lower methylation levels in ecDNA+ samples. We calculated the methylation levels of the coamplified fragments in ecDNA+ samples (ecDNA+ coamplification regions), the non-coamplified fragments in ecDNA+ samples (ecDNA+ no coamplification regions), and the corresponding linear chromosome regions in ecDNA- samples (ecDNA- chromosome regions) in above 14 tumors. The results showed (Figure 4C, Supplementary Table 4) that in most tumors (12 out of 14), the methylation levels were lower in ecDNA+ samples with co-amplification compared to ecDNA- samples. The most significant differences were found in CESC, ESCA, and LUSC, with p-values <1e-16. Notably, in 10 tumors (BLCA, BRCA, CESC, COAD, ESCA, LGG, LUAD, LUSC, SKCM, and STAD), the methylation levels were lower in ecDNA+ samples with co-amplification than in ecDNA+ samples without co-amplification. Among them, CESC, ESCA, and SKCM showed the highest significance, with p-values <2.22e-16. It is indicated that the lower methylation levels on ecDNA are common in tumors, and this low methylation tendency is more likely to occur in ecDNA+ samples with co-amplification.

### 3.5. ecDNA Regions Exhibit Higher Accessibility and Recruit Transcription Factors Involved in Tumor Biology Processes

Previous studies have shown that DNA fragments circularized into ecDNA exhibit higher accessibility and can influence the transcription and expression of oncogenes [7][22]. Therefore, we integrated TCGA ATAC data to identify accessible peaks in 6 tumors (BLCA, BRCA, UCEC, ESCA, SKCM, and STAD). We then compared the genome-wide accessibility levels between ecDNA+ and ecDNA- samples in these 6 tumors to investigate whether the presence of ecDNA in patients affects the overall genome accessibility. The results showed that only BLCA, BRCA, and UCEC exhibited significant differences (Supplementary Figure S10), indicating that the presence or absence of ecDNA is not the key determinant of genome accessibility. Moreover, we analyzed the accessibility of ecDNA regions in the 6 tumor types compared to the corresponding linear regions in ecDNA- samples. The results revealed significantly higher accessibility in ecDNA regions compared to the corresponding linear regions in all 6 tumors (Figure 5A). The overall accessibility of ecDNA regions exhibits a certain degree of chromosome preference, as in most tumors, the amplified fragments from ecDNA in chr3, 6, 15, and 20 exhibited more enriched ATAC peaks compared to other chromosomes (Figure 5B).

**Figure 5.**
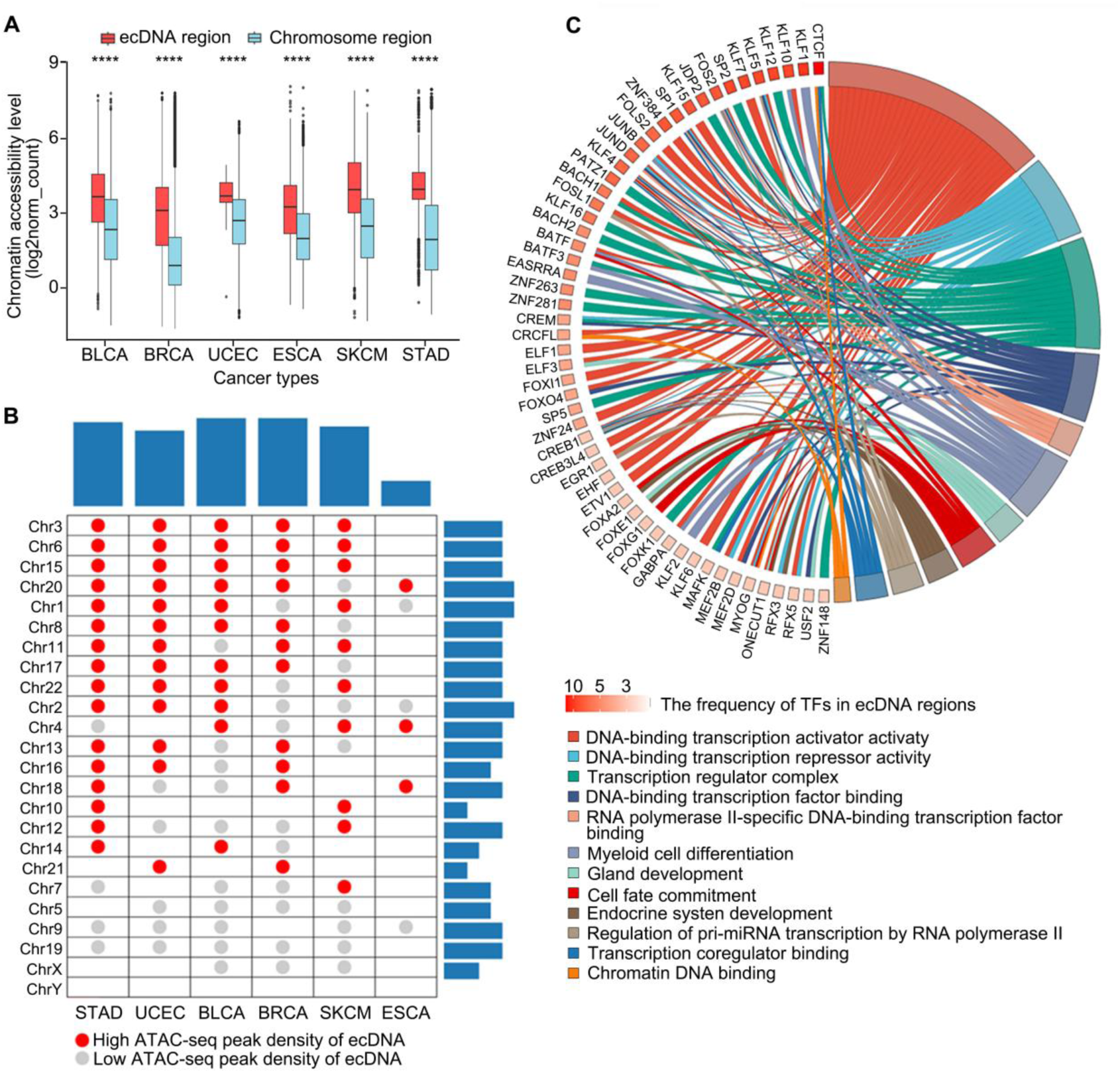
Analysis of accessibility and transcription factors binding in ecDNA. A. Accessibility of ecDNA regions compared to the corresponding linear regions in 6 tumors. The significance was determined using the paired Wilcoxon test (*p≤0.05, **p≤0.01, ***p≤0.001, nsp > 0.05). B. Analysis of accessibility preferences of ecDNA-originating chromosomes in 6 tumors. The top bar graph indicates the number of accessible chromosomes for each tumor’s ecDNA origin, while the right bar graph represents the number of tumors in which ecDNA originates from each chromosome. C. GO enrichment analysis of the 201 transcription factors found in ecDNA regions. The gradient color indicates the occurrence frequency of each transcription factor in ecDNA.

Transcription factors play an important role in chromatin accessibility-mediated transcriptional regulation. Therefore, we integrated a transcription factor database to identify 201 significantly enriched transcription factors in ecDNA accessible regions (Supplementary Table 5). Transcription factors that are frequently enriched in multiple ecDNA regions often indicate their important functions in ecDNA. Therefore, we conducted GO analysis to examine the biological processes involved in these 201 transcription factors. We found that the top 10 transcription factors frequently occurring in ecDNA regions are mainly associated with transcriptional regulation, expression regulation, DNA binding, and tumor development. Among them, CTCF, KLF family, and SP family are primarily enriched in transcriptional regulation processes such as DNA-binding transcription activator activity and transcription regulator complex (Figure 5C). For instance, it has been previously reported that CTCF can serve as a marker for TAD boundaries, and the formation of new TADs in ecDNA requires convergent directionality of CTCF sites to establish a new boundary and drive abnormal gene expression [23]. Taken together, these results suggested that ecDNA regions exhibit significantly higher accessibility levels and present a certain degree of chromosome preferences. Transcription factors that frequently bind to ecDNA are commonly involved in ecDNA- related transcriptional regulatory processes.

### 3.6. ecDNA Significantly Influences the Expression and Pathways of Tumor Antigen P resentation Genes

Tumor immunity refers to the process by which the immune system combats tumor development and suppresses tumor growth. However, tumor progression can evade immune clearance through mechanisms such as immune escape, leading to further deterioration of the tumor. To investigate whether ecDNA regulates immune checkpoints and induces immune escape, we examined the expression of 13 common immune checkpoint genes in ecDNA+ and ecDNA- samples (Supplementary Figure S11). The results showed that, except for B7-H4, most immune checkpoint genes did not exhibit significantly higher expression in ecDNA+ samples. This suggested that immune escape in ecDNA+ tumors is not primarily mediated through the stimulation of immune checkpoint signaling. Genes related to antigen presentation are responsible for presenting tumor-derived antigens to the immune system, triggering an immune response. However, tumor cells can reduce the effectiveness of antigen presentation to evade immune surveillance and attack, leading to immune escape. To investigate whether ecDNA regulates genes involved in antigen presentation and leads to immune escape, we analyzed the expression of antigen presentation-related genes in ecDNA+ and ecDNA- samples. The results showed that most antigen presentation genes were downregulated in ecDNA+ samples (Figure 6A). Among them, HSPH1, TLR4, and HLA-DOA exhibited significantly lower expression in ecDNA+ samples, with p-values of 8.8e-06, 4.0e-05, and 4.99e-05, respectively. The downregulation of antigen presentation genes in ecDNA+ patients suggested a restriction in the efficient presentation of antigens to the immune system, leading to a reduced ability of the immune system to recognize and respond to antigens [17]. To further investigate whether the pathways associated with antigen presentation are also downregulated in ecDNA+ samples, we calculated the activity levels of 4 antigen presentation-related pathways using the R package “GSVA”. As shown in Figure 6B, the GSVA scores of antigen presentation pathways corresponding to ecDNA+ samples were significantly lower compared to those corresponding to ecDNA- samples. This suggested that ecDNA+ tumors may inhibit immune responses in ecDNA+ patients by downregulating antigen presentation pathways and limiting effective antigen presentation.

**Figure 6.**
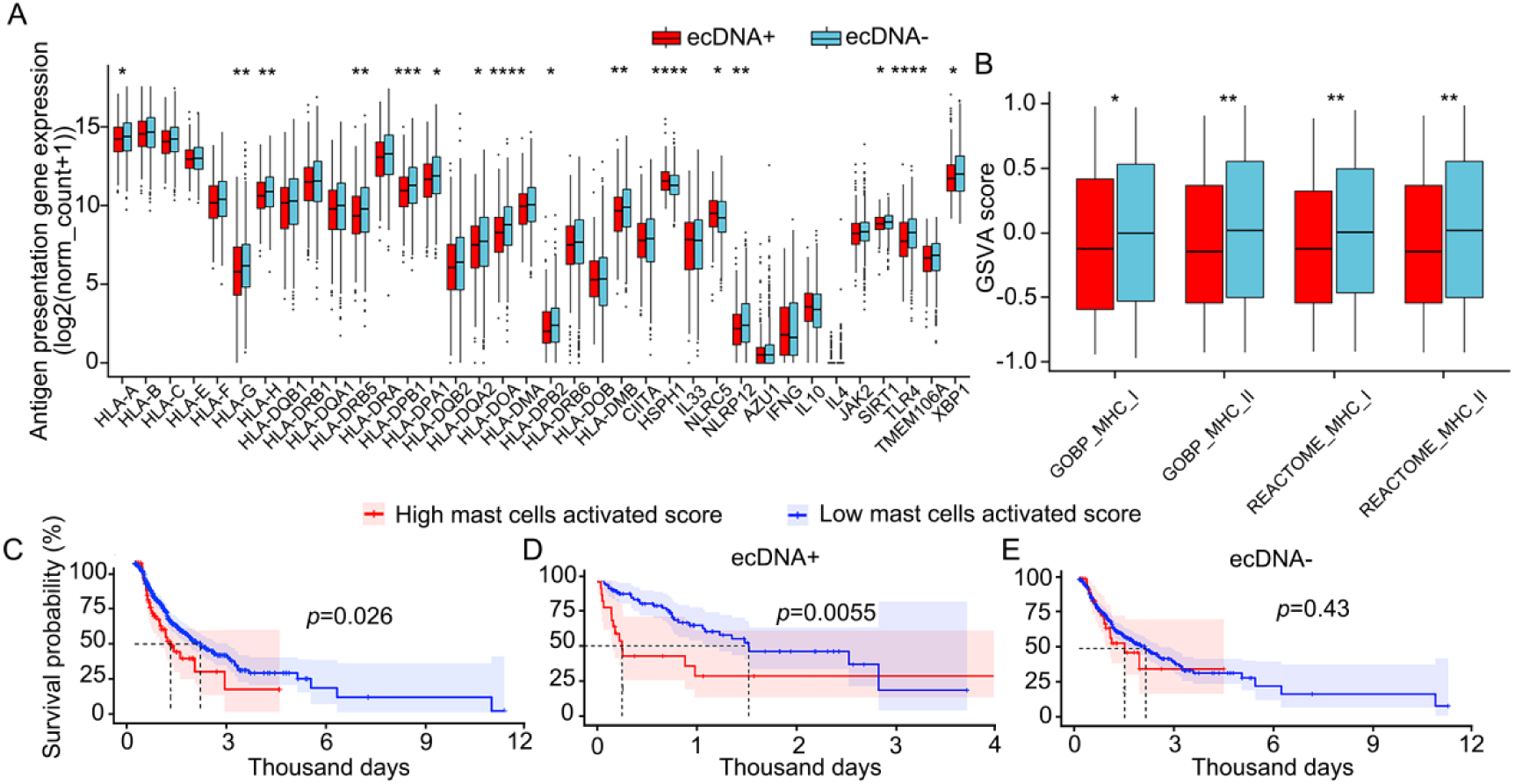
Analysis of antigen presentation and immune cell infiltration in ecDNA. A. Expression of antigen presentation genes in ecDNA+ and ecDNA- samples. The significance was determined using the Wilcoxon test (*p≤0.05, **p≤0.01, ***p≤0.001, nsp>0.05). B. Activity levels of antigen presentation-related pathways in ecDNA+ and ecDNA- samples; higher GSVA score indicates higher activity levels of antigen presentation-related genes, while lower GSVA score indicates lower activity levels of antigen presentation-related genes. C. Degree of mast cell activation and patient survival in tumors. D. Impact of mast cell infiltration on the prognosis and survival of ecDNA+ patients. E. Impact of mast cell infiltration on the prognosis and survival of ecDNA- patients.

Quantification of immune cell infiltration in tumors helps reveal the multifaceted role of the immune system in human cancer, including its involvement in tumor escape mechanisms and response to treatment, thus facilitating the understanding of disease occurrence and progression. We then investigated the infiltration levels of 22 immune cell types in ecDNA+ and ecDNA- patients (Supplementary Figure S12). The results showed significantly higher levels of activated mast cells in ecDNA+ samples (p<0.01). Mast cell activation has been reported as a potential mediator of cancer-related malignancies and is associated with tumor development [24], suggesting that activated mast cells may play a significant role in immune escape in ecDNA+ patients. Furthermore, we utilized LASSO and COX analyses to assess the impact of mast cell activation on patient survival. The results demonstrated a significant association between the extent of mast cell activation and patient survival (Figure 6C). In addition, we investigated the influence of activated mast cells on the prognosis of ecDNA+ and ecDNA- patients. In ecDNA+ patients, a higher degree of infiltration of activated mast cells significantly shortened patient survival compared to a lower degree of infiltration of activated mast cells (Figure 6D). In contrast, the infiltration level of activated mast cells did not affect patient survival in ecDNA- patients (Figure 6E). Taken together, these results suggested that the downregulation of antigen presentation gene expression and antigen presentation pathway leads to immune escape in ecDNA tumors. Moreover, the significant infiltration of activated mast cells in ecDNA+ tumors is associated with poorer prognosis in ecDNA+ patients.

---

**Supplementary Fig. S1.**
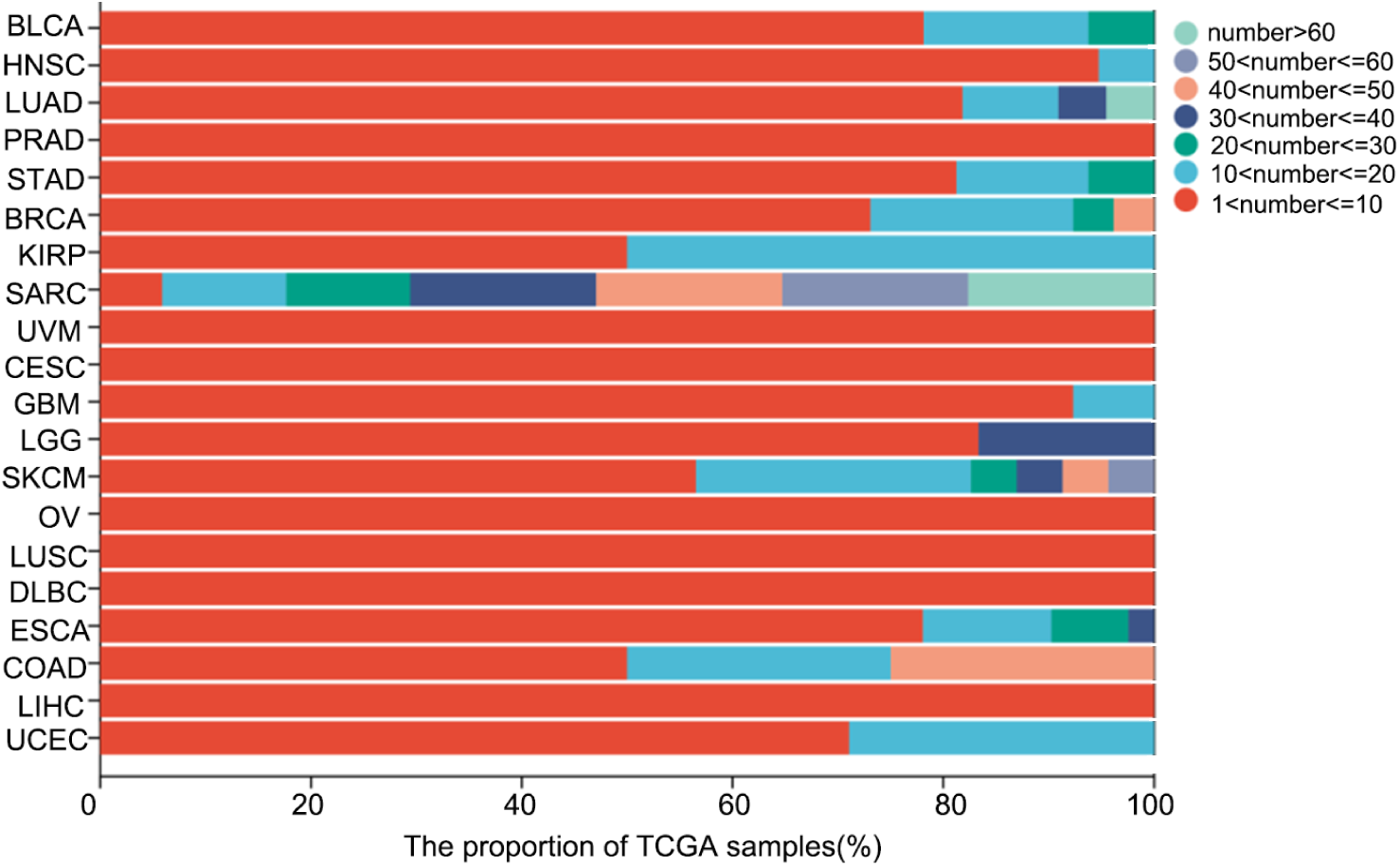
The proportion of TCGA samples in 20 tumors.

**Supplementary Fig. S2.**
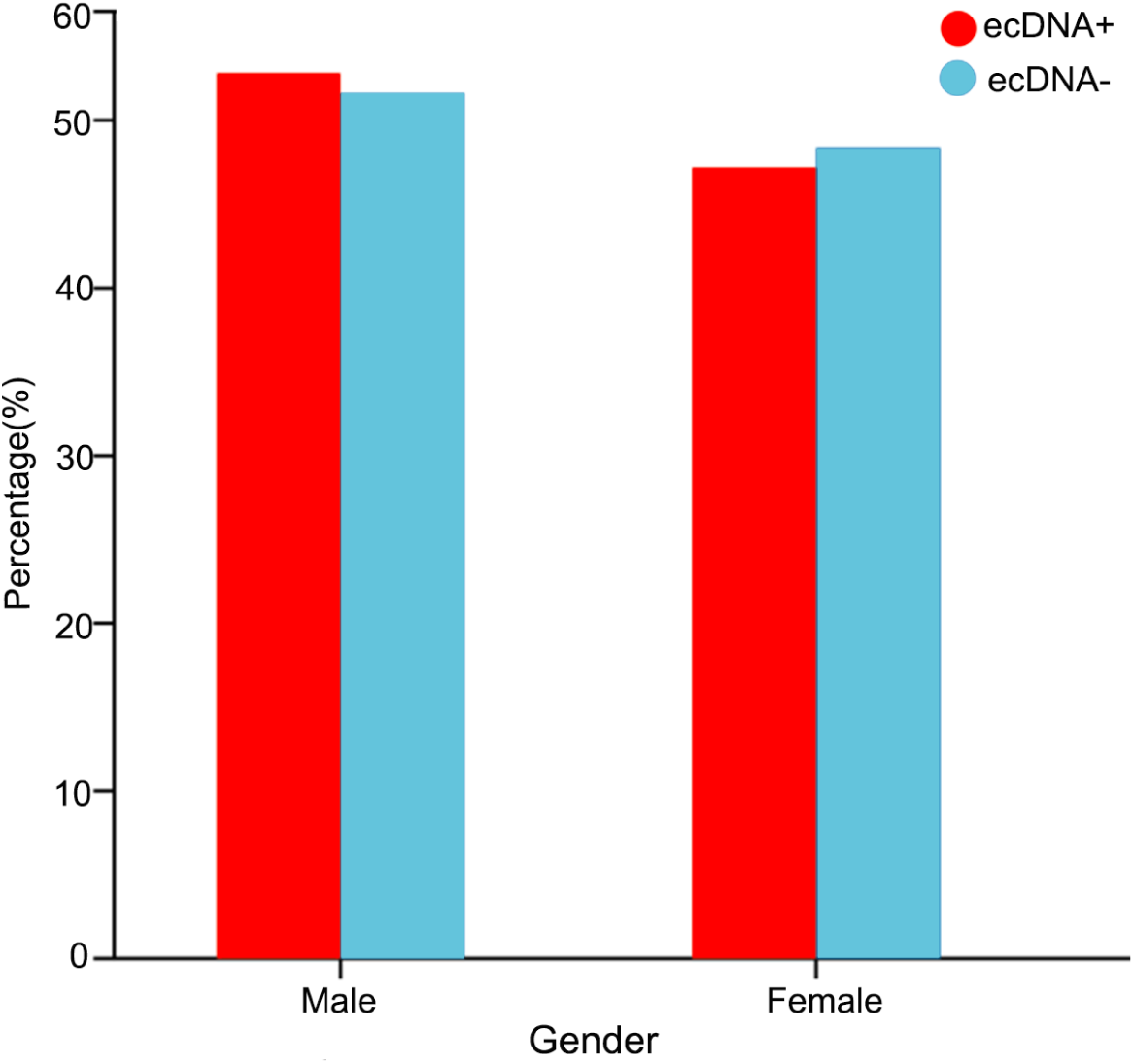
The gender distribution of ecDNA+ and ecDNA- samples in 20 tumors.

**Supplementary Fig. S3.**
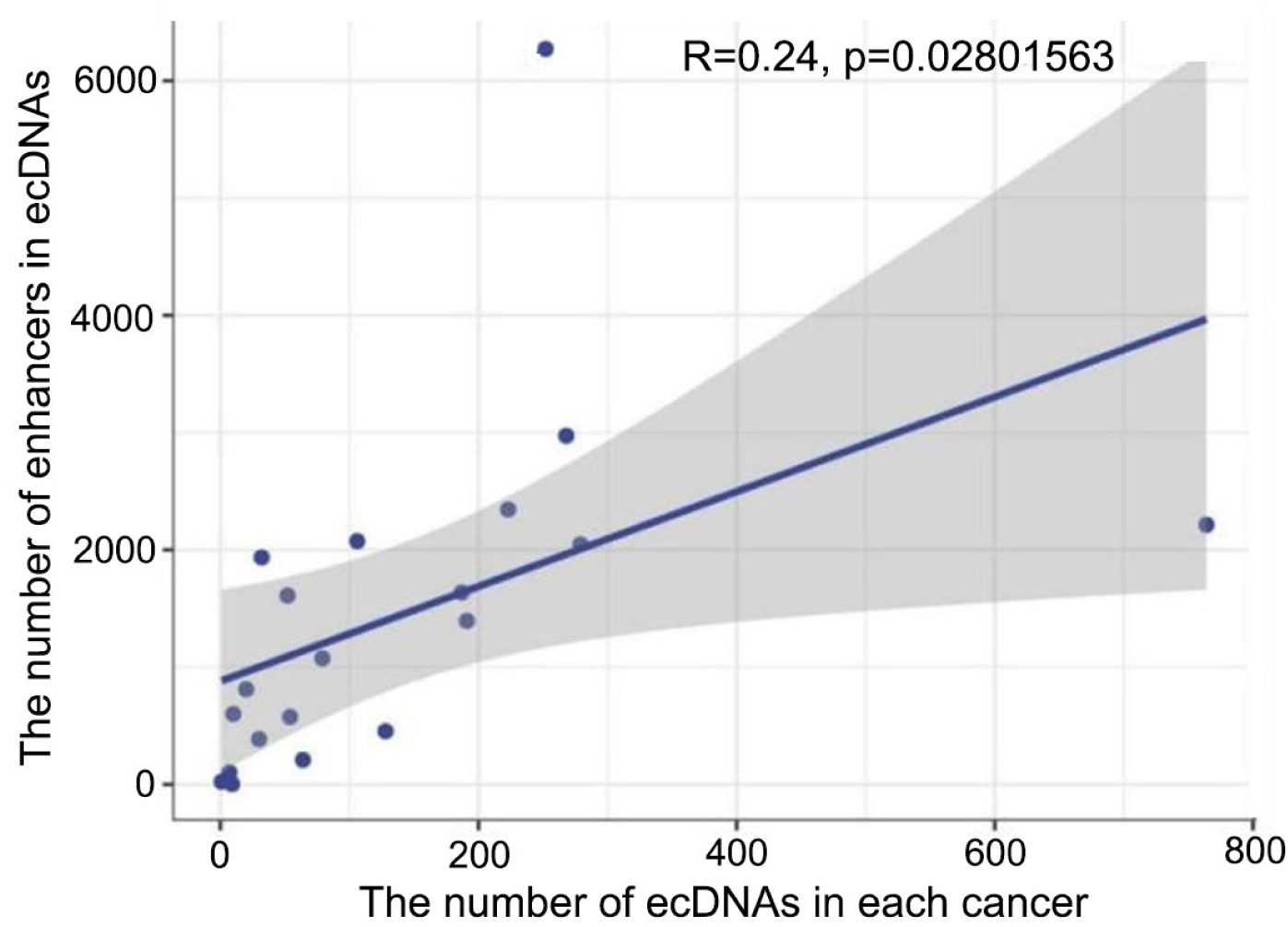
The correlation between the total number of ecDNAs and enhancers; The plot represents a comprehensive view of 20 tumors, with a total of 20 data points, each corresponding to a specific cancer type.

**Supplementary Fig. S4.**
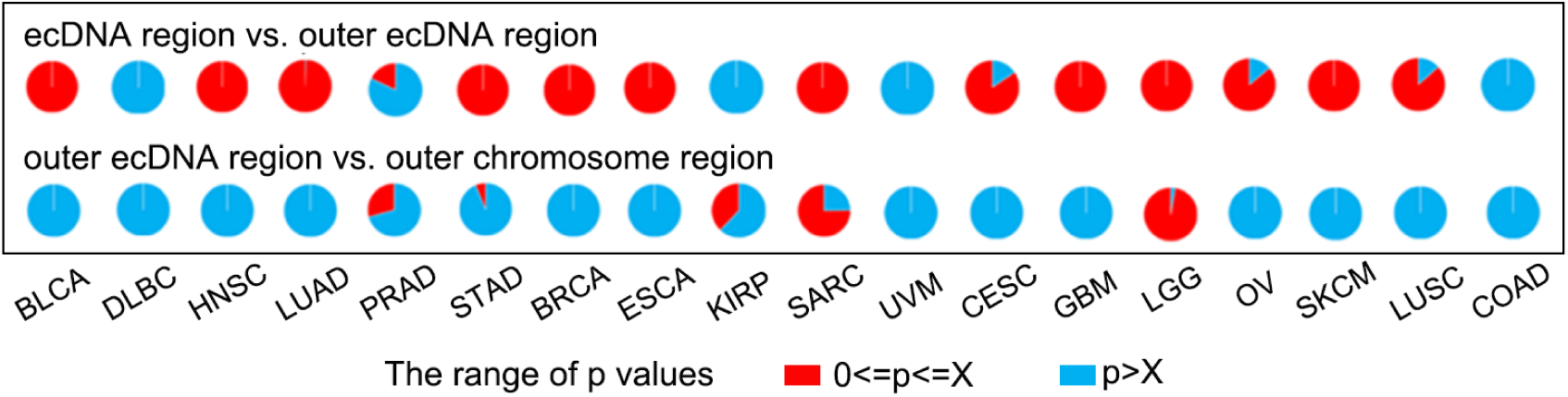
The eRNA expression in ecDNA regions and the corresponding linear chromosome regions was randomly sampled in 18 cancers, along with the expression in regions outside of ecDNA. The value ‘X’ represents the P-value of eRNA expression in the ecDNA regions and the corresponding linear chromosome regions in each cancer. The first row of pie graphs shows the percentage of significant p-values for enhancer expression in ecDNA regions and regions outside of ecDNA. The blue color (p>X) indicates that enhancers have lower expression levels in the ecDNA regions compared to the regions outside of ecDNA, while the red color (0<=p<=X) indicates that enhancers have higher expression levels in the ecDNA regions compared to the regions outside of ecDNA. The second row of pie graphs shows the percentage of significant p-values for enhancer expression in regions outside of ecDNA and the corresponding linear chromosome regions. The blue color (p>X) indicates that enhancers have higher expression levels in regions outside of ecDNA compared to the corresponding linear chromosome regions, while the red color (0<=p<=X) indicates that enhancers have lower expression levels in regions outside of ecDNA compared to the corresponding linear chromosome regions.

**Supplementary Fig. S5.**
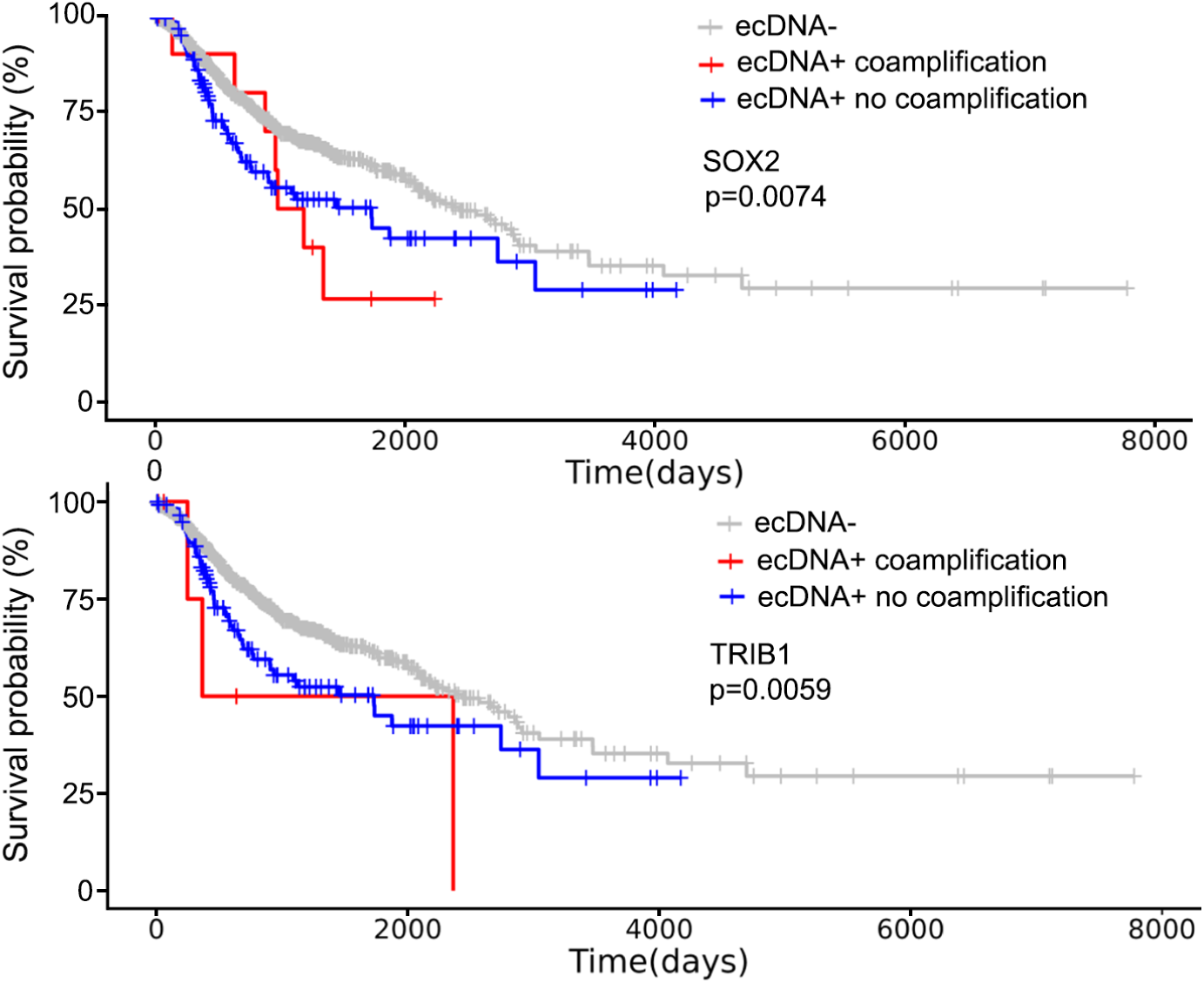
The impact of SOX2 and TRIB1 on the prognosis of ecDNA-, ecDNA+ with co-amplification, and ecDNA+ without co-amplification patients.

**Supplementary Fig. S6.**
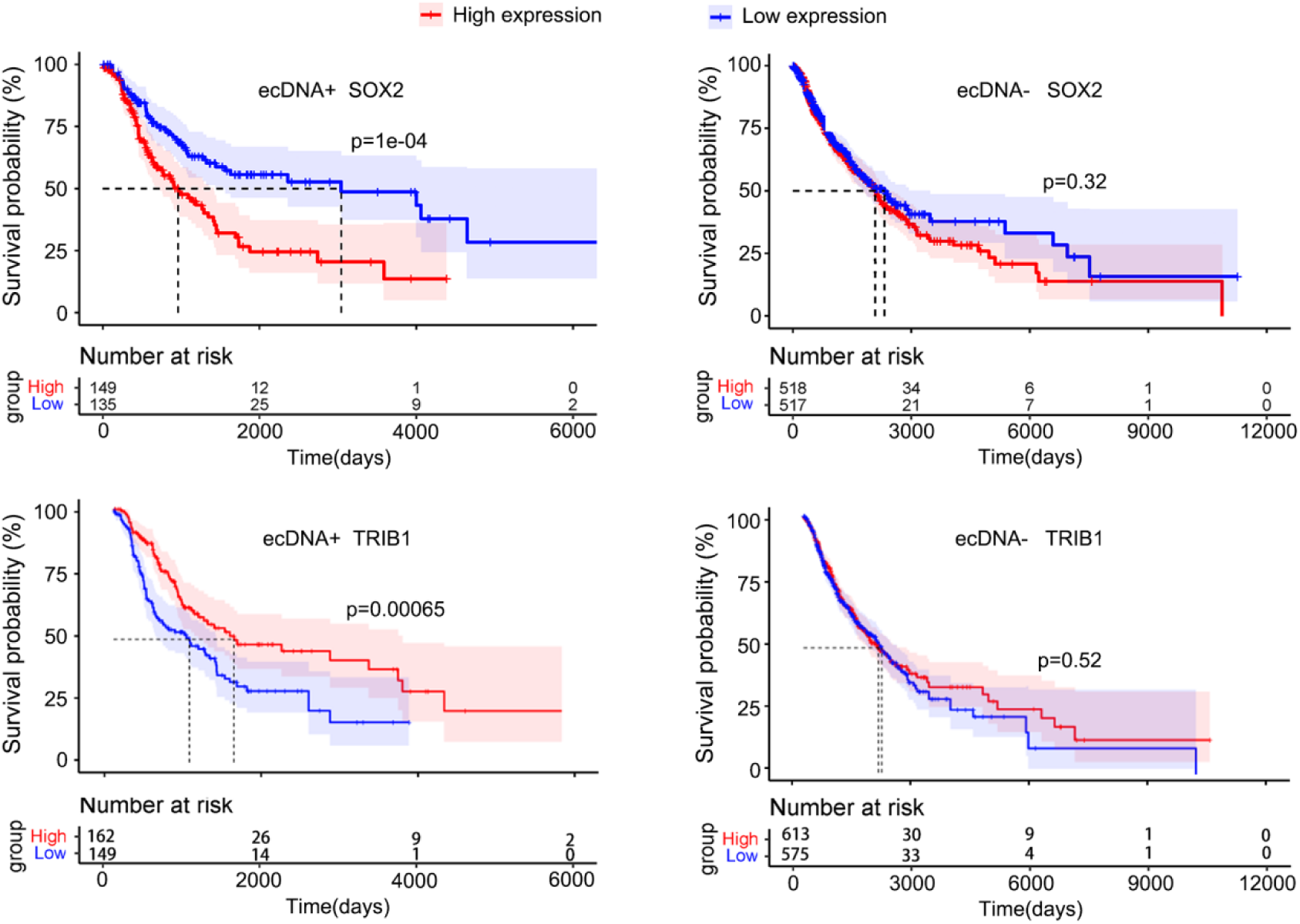
The survival curves of ecDNA+ and ecDNA- patients in relation to SOX2 and TRIB1.

**Supplementary Fig. S7.**
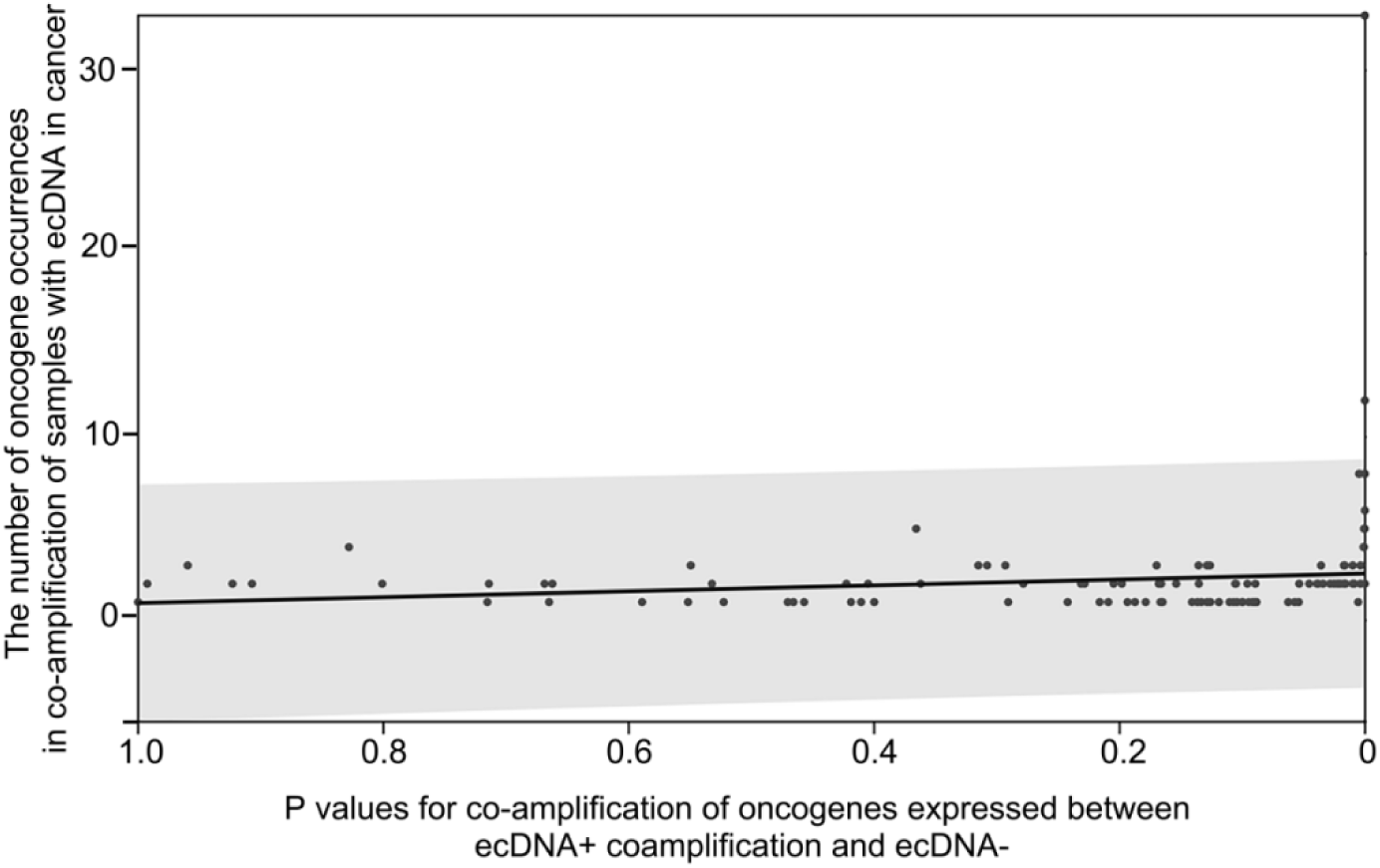
The Spearman correlation between the frequency and the expression levels of co-amplified oncogenes in ecDNA+ with co-amplification and ecDNA- samples, as indicated by the p-values. A higher number of oncogenes involved in co-amplification may be associated with a more significant difference in expression levels between ecDNA+ with co-amplification and ecDNA- samples.

**Supplementary Fig. S8.**
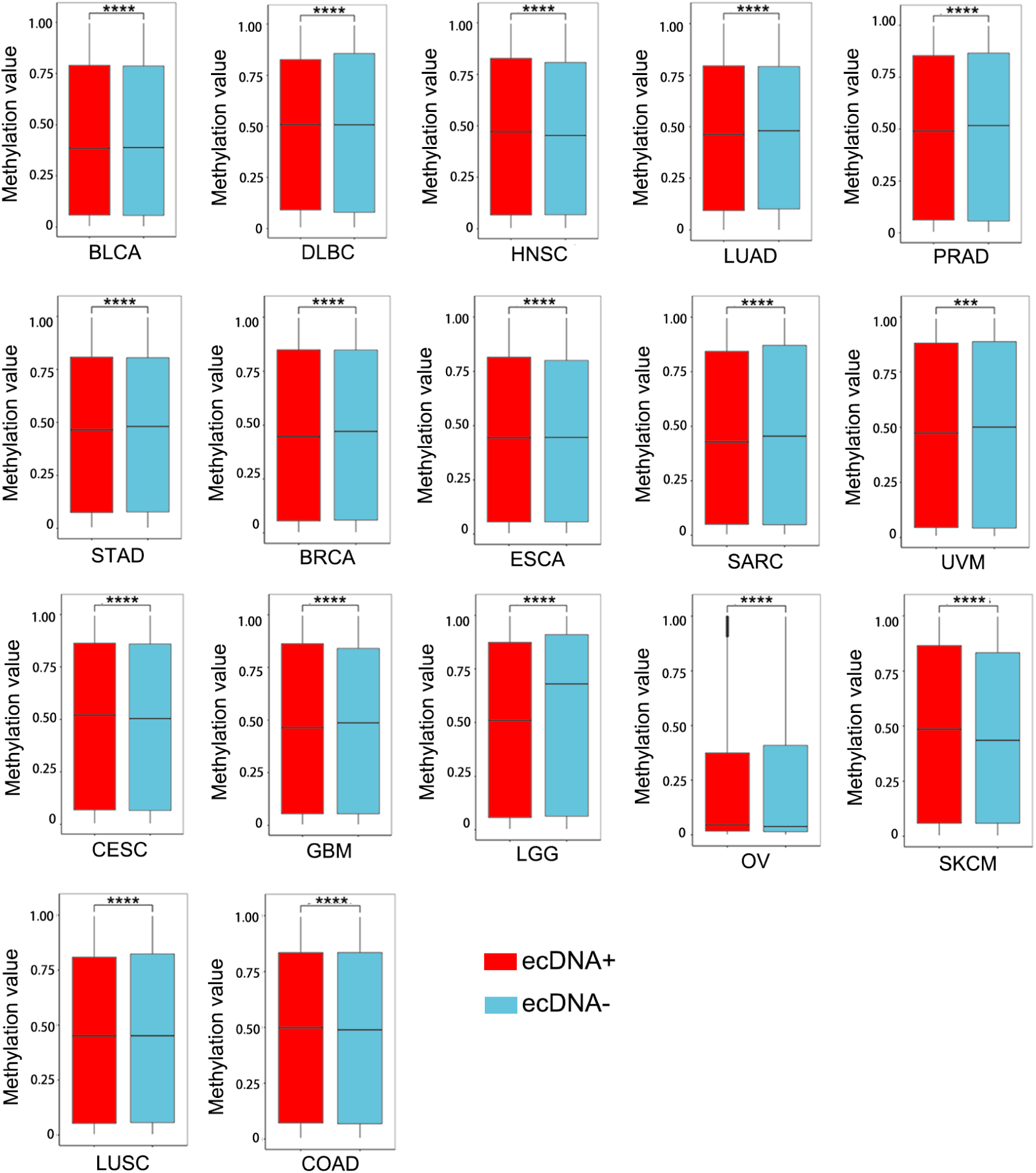
Comparison of methylation levels between ecDNA+ and ecDNA- samples in 17 cancers. The red color corresponds to ecDNA+ samples, while the blue color corresponds to ecDNA- samples. In the cases of BLCA, LUAD, PRAD, STAD, BRCA, SARC, UVM, GBM, LGG, and LUSC, the methylation levels in ecDNA+ samples are significantly lower than in ecDNA- samples. In contrast, in the cases of DLBC, HNSC, ESCA, CESC, GBM, SKCM, and COAD, the methylation levels in ecDNA+ samples are significantly higher than in ecDNA- samples. The significance was determined using a Wilcoxon test (*p≤0.05, **p≤0.01, ***p≤0.001, ^ns^p>0.05).

**Supplementary Fig. S9.**
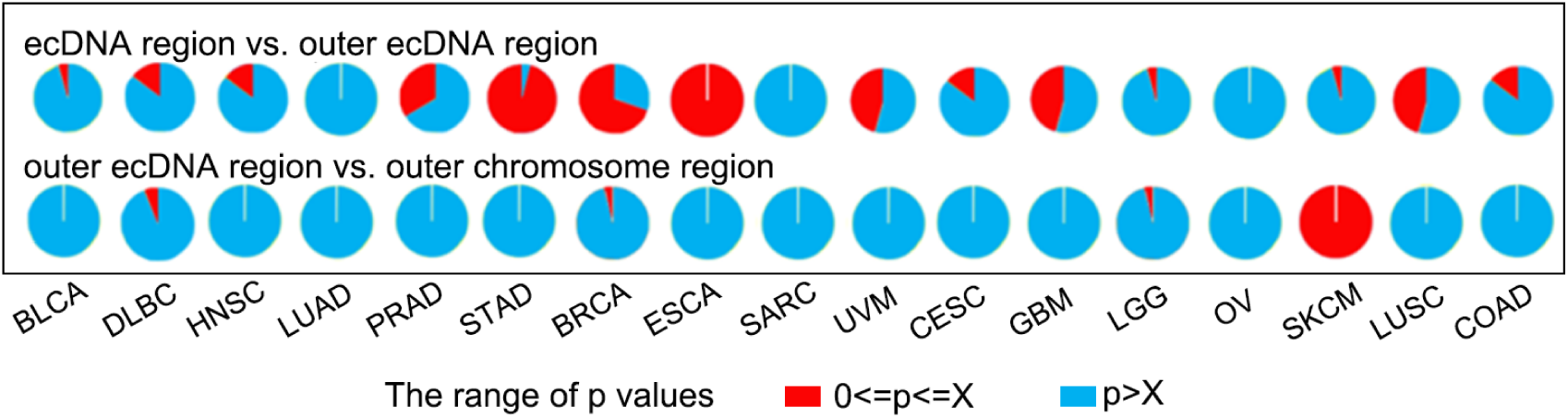
Under the conditions of random sampling in 17 tumors, the proportions of significantly different methylation levels are represented using pie graphs; The first row of pie graphs illustrates the percentage of significant p-values for methylation in the ecDNA regions and the regions outside of ecDNA. The blue color (p>X) indicates that methylation does not occur in the ecDNA regions, while the red color (0<=p<=X) indicates that methylation occurs in the ecDNA regions. The second row of pie graphs displays the percentage of significant p-values for methylation in regions outside of ecDNA and the corresponding linear chromosome regions. The blue color (p>X) suggests that methylation occurs in the ecDNA regions, whereas the red color (0<=p<=X) suggests that methylation occurs outside of the ecDNA regions; X represents the P values of methylation level in ecDNA regions and the corresponding linear chromosome regions.

**Supplementary Fig. S10.**
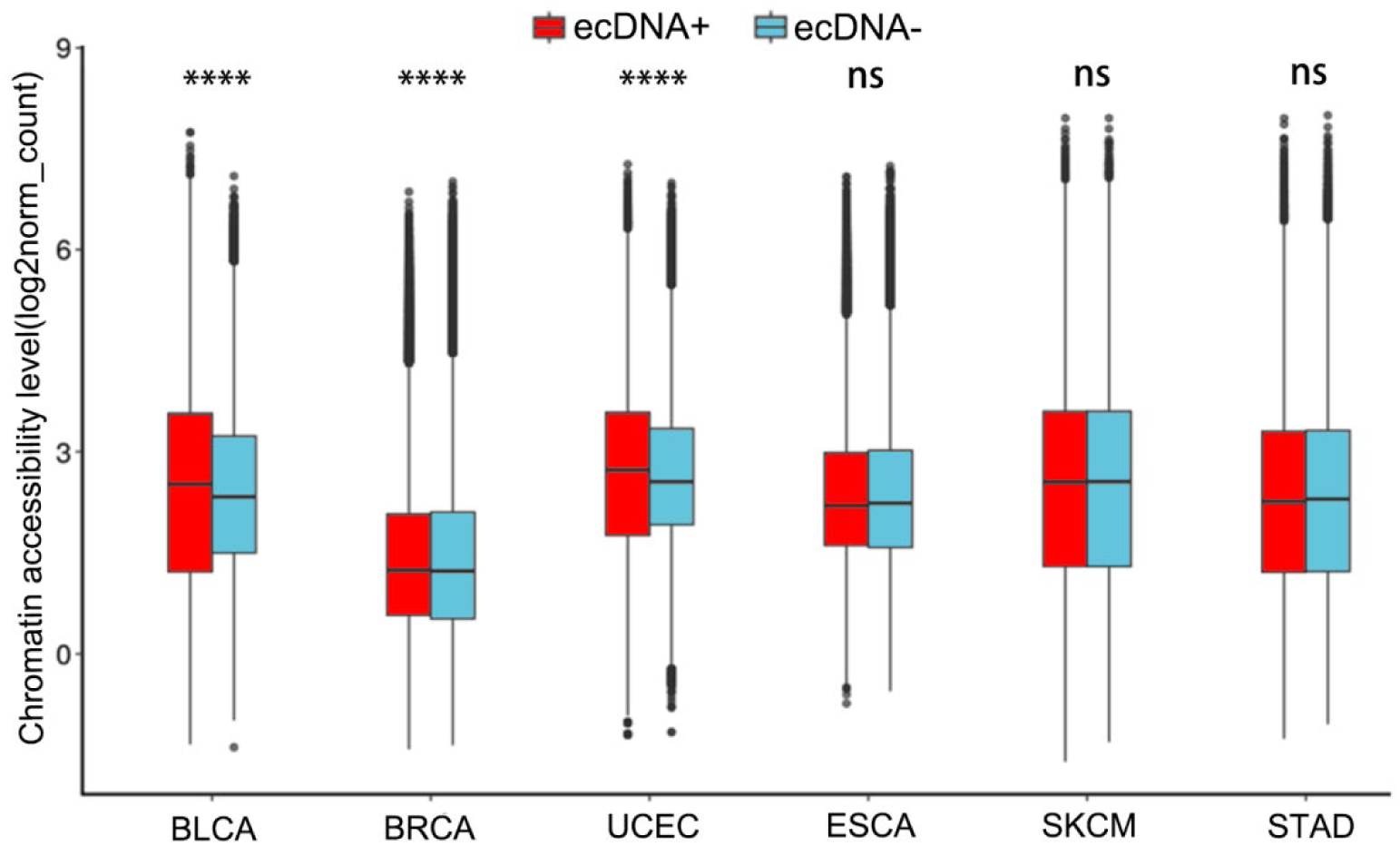
The chromatin accessibility level of ecDNA+ and ecDNA- samples in 6 tumors. The significance was determined using a Wilcoxon test. (*p≤0.05, **p≤0.01, ***p≤0.001, ^ns^p>0.05).

**Supplementary Fig. S11.**
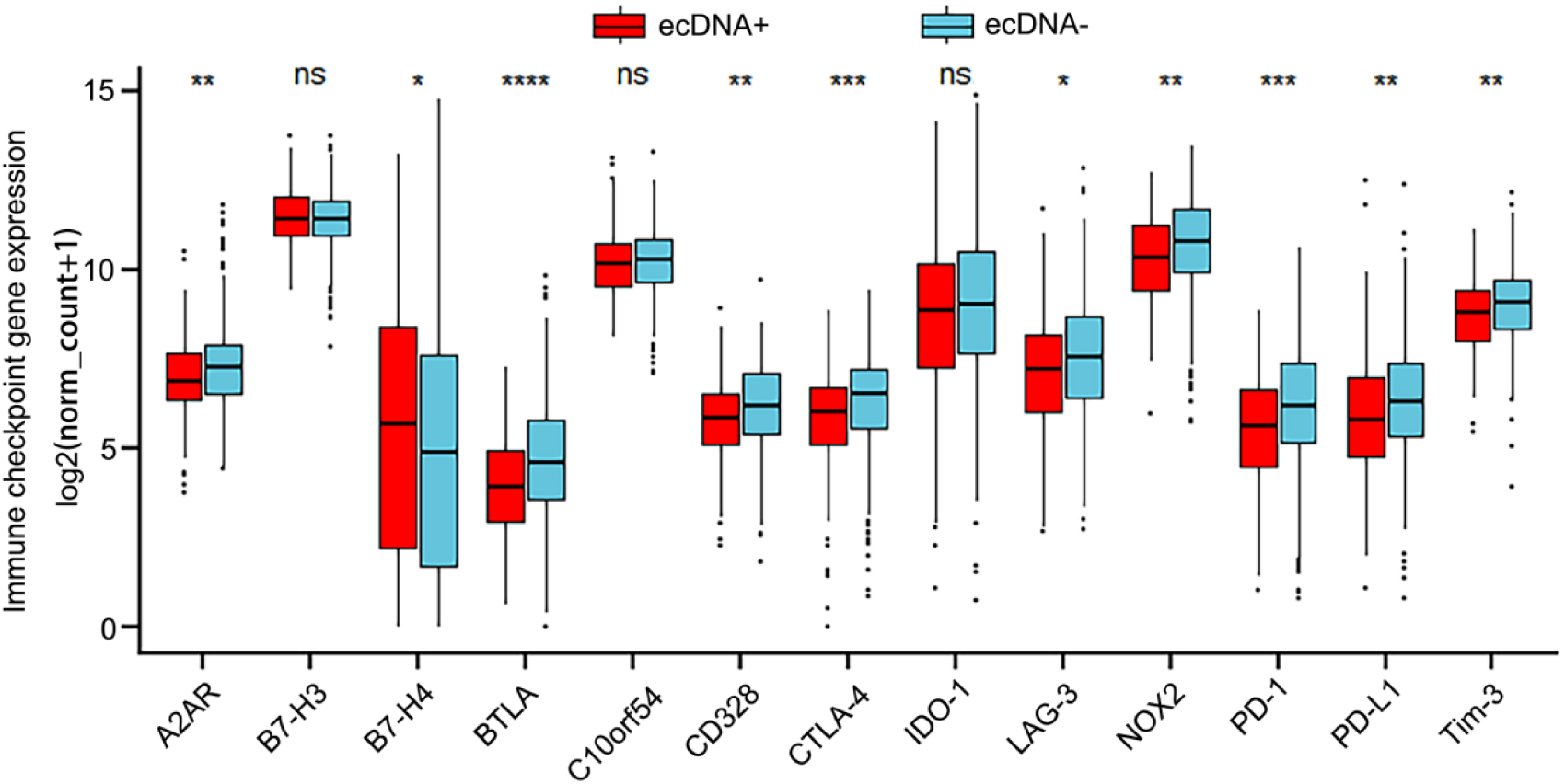
The expression of immune checkpoint genes in ecDNA+ and ecDNA- samples. The significance was determined using a Wilcoxon test. (*p≤0.05, **p≤0.01, ***p≤0.001, ^ns^p>0.05).

**Supplementary Fig. S12.**
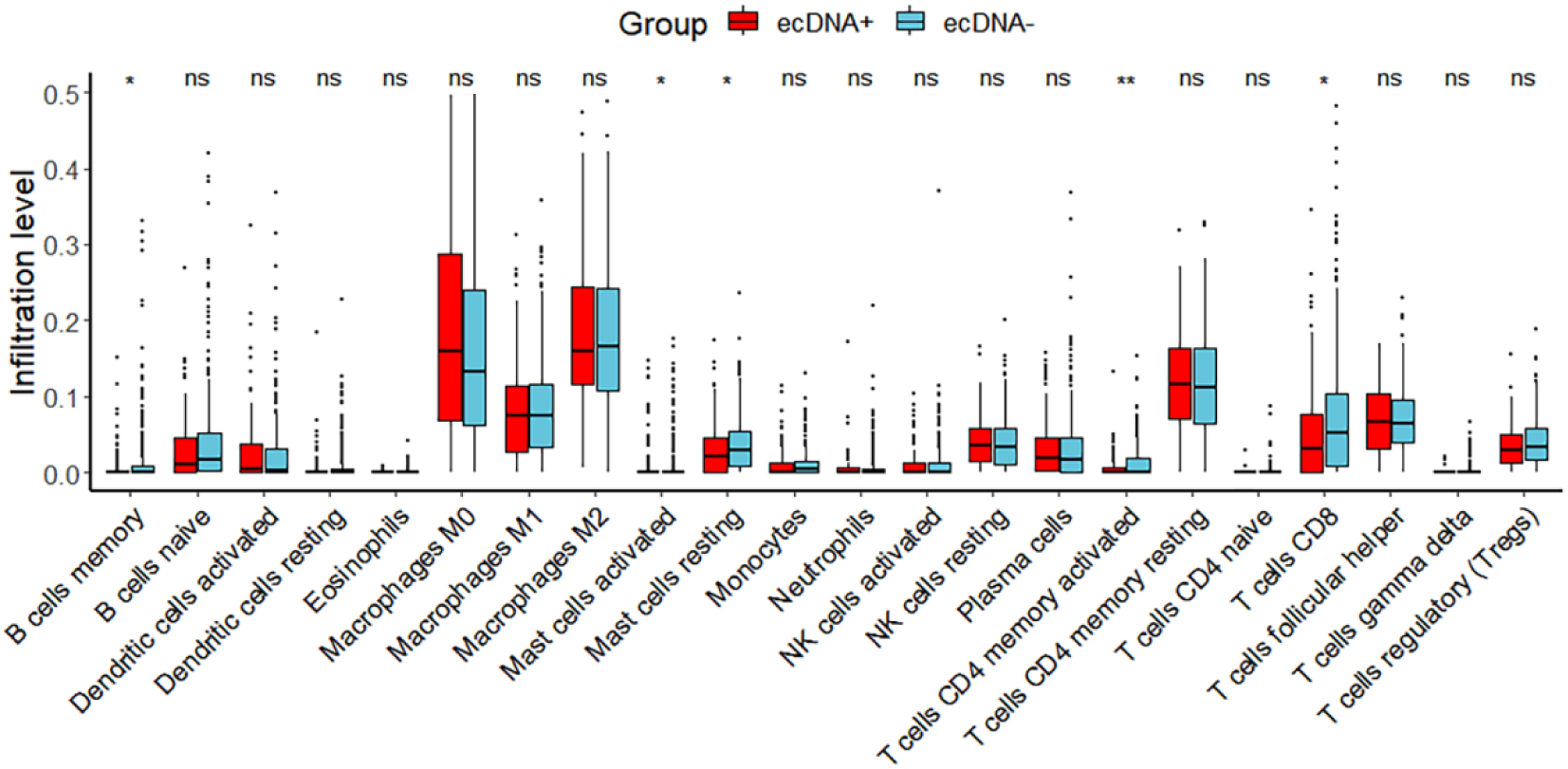
The infiltration of 22 types of immune cells in ecDNA+ and ecDNA- samples. The significance was determined using a Wilcoxon test. (*p≤0.05, **p≤0.01, ***p≤0.001, ^ns^p>0.05).

## References

[1] Turner, K. M., Deshpande, V., Beyter, D., Koga, T., Rusert, J., Lee, C., Li, B., Arden, K., Ren, B., Nathanson, D. A., Kornblum, H. I., Taylor, M. D., Kaushal, S., Cavenee, W. K., Wechsler-Reya, R., Furnari, F. B., Vandenberg, S. R., Rao, P. N., Wahl, G. M., Bafna, V., … Mischel, P. S. (2017). Extrachromosomal oncogene amplification drives tumour evolution and genetic heterogeneity. Nature, 543(7643), 122–125. 10.1038/nature21356

[2] Kim, H., Nguyen, N. P., Turner, K., Wu, S., Gujar, A. D., Luebeck, J., Liu, J., Deshpande, V., Rajkumar, U., Namburi, S., Amin, S. B., Yi, E., Menghi, F., Schulte, J. H., Henssen, A. G., Chang, H. Y., Beck, C. R., Mischel, P. S., Bafna, V., & Verhaak, R. G. W. (2020). Extrachromosomal DNA is associated with oncogene amplification and poor outcome across multiple cancers. Nature genetics, 52(9), 891–897. 10.1038/s41588-020-0678-2

[3] Andersson, R., Gebhard, C., Miguel-Escalada, I., Hoof, I., Bornholdt, J., Boyd, M., Chen, Y., Zhao, X., Schmidl, C., Suzuki, T., Ntini, E., Arner, E., Valen, E., Li, K., Schwarzfischer, L., Glatz, D., Raithel, J., Lilje, B., Rapin, N., Bagger, F. O., … Sandelin, A. (2014). An atlas of active enhancers across human cell types and tissues. Nature, 507(7493), 455–461. 10.1038/nature12787

[4] Li, Q., Peterson, K. R., Fang, X., & Stamatoyannopoulos, G. (2002). Locus control regions. Blood, 100(9), 3077–3086. 10.1182/blood-2002-04-1104

[5] Morton, A. R., Dogan-Artun, N., Faber, Z. J., MacLeod, G., Bartels, C. F., Piazza, M. S., Allan, K. C., Mack, S. C., Wang, X., Gimple, R. C., Wu, Q., Rubin, B. P., Shetty, S., Angers, S., Dirks, P. B., Sallari, R. C., Lupien, M., Rich, J. N., & Scacheri, P. C. (2019). Functional Enhancers Shape Extrachromosomal Oncogene Amplifications. Cell, 179(6), 1330–1341.e13. 10.1016/j.cell.2019.10.039

[6] Helmsauer, K., Valieva, M. E., Ali, S., Chamorro González, R., Schöpflin, R., Röefzaad, C., Bei, Y., Dorado Garcia, H., Rodriguez-Fos, E., Puiggròs, M., Kasack, K., Haase, K., Keskeny, C., Chen, C. Y., Kuschel, L. P., Euskirchen, P., Heinrich, V., Robson, M. I., Rosswog, C., Toedling, J., … Koche, R. P. (2020). Enhancer hijacking determines extrachromosomal circular MYCN amplicon architecture in neuroblastoma. Nature communications, 11(1), 5823. 10.1038/s41467-020-19452-y

[7] Zhu, Y., Gujar, A. D., Wong, C. H., Tjong, H., Ngan, C. Y., Gong, L., Chen, Y. A., Kim, H., Liu, J., Li, M., Mil-Homens, A., Maurya, R., Kuhlberg, C., Sun, F., Yi, E., deCarvalho, A. C., Ruan, Y., Verhaak, R. G. W., & Wei, C. L. (2021). Oncogenic extrachromosomal DNA functions as mobile enhancers to globally amplify chromosomal transcription. Cancer cell, 39(5), 694–707.e7. 10.1016/j.ccell.2021.03.006

[8] Hung, K. L., Yost, K. E., Xie, L., Shi, Q., Helmsauer, K., Luebeck, J., Schöpflin, R., Lange, J. T., Chamorro González, R., Weiser, N. E., Chen, C., Valieva, M. E., Wong, I. T., Wu, S., Dehkordi, S. R., Duffy, C. V., Kraft, K., Tang, J., Belk, J. A., Rose, J. C., … Chang, H. Y. (2021). ecDNA hubs drive cooperative intermolecular oncogene expression. Nature, 600(7890), 731–736. 10.1038/s41586-021-04116-8

[9] Jiang, X., Pan, X., Li, W., Han, P., Yu, J., Li, J., Zhang, H., Lv, W., Zhang, Y., He, Y., & Xiang, X. (2023). Genome-wide characterization of extrachromosomal circular DNA in gastric cancer and its potential role in carcinogenesis and cancer progression. Cellular and molecular life sciences : CMLS, 80(7), 191. 10.1007/s00018-023-04838-0

[10] Hung, K. L., Luebeck, J., Dehkordi, S. R., Colón, C. I., Li, R., Wong, I. T., Coruh, C., Dharanipragada, P., Lomeli, S. H., Weiser, N. E., Moriceau, G., Zhang, X., Bailey, C., Houlahan, K. E., Yang, W., González, R. C., Swanton, C., Curtis, C., Jamal-Hanjani, M., Henssen, A. G., … Chang, H. Y. (2022). Targeted profiling of human extrachromosomal DNA by CRISPR-CATCH. Nature genetics, 54(11), 1746–1754. 10.1038/s41588-022-01190-0

[11] Yi, E., Gujar, A. D., Guthrie, M., Kim, H., Zhao, D., Johnson, K. C., Amin, S. B., Costa, M. L., Yu, Q., Das, S., Jillette, N., Clow, P. A., Cheng, A. W., & Verhaak, R. G. W. (2022). Live-Cell Imaging Shows Uneven Segregation of Extrachromosomal DNA Elements and Transcriptionally Active Extrachromosomal DNA Hubs in Cancer. Cancer discovery, 12(2), 468–483. 10.1158/2159-8290.CD-21-1376

[12] Pongor, L. S., Schultz, C. W., Rinaldi, L., Wangsa, D., Redon, C. E., Takahashi, N., Fialkoff, G., Desai, P., Zhang, Y., Burkett, S., Hermoni, N., Vilk, N., Gutin, J., Gergely, R., Zhao, Y., Nichols, S., Vilimas, R., Sciuto, L., Graham, C., Caravaca, J. M., … Thomas, A. (2023). Extrachromosomal DNA Amplification Contributes to Small Cell Lung Cancer Heterogeneity and Is Associated with Worse Outcomes. Cancer discovery, 13(4), 928–949. 10.1158/2159-8290.CD-22-0796

[13] Prada-Luengo, I., Krogh, A., Maretty, L., & Regenberg, B. (2019). Sensitive detection of circular DNAs at single-nucleotide resolution using guided realignment of partially aligned reads. BMC bioinformatics, 20(1), 663. 10.1186/s12859-019-3160-3

[14] Chen, H., Li, C., Peng, X., Zhou, Z., Weinstein, J. N., Cancer Genome Atlas Research Network, & Liang, H. (2018). A Pan-Cancer Analysis of Enhancer Expression in Nearly 9000 Patient Samples. Cell, 173(2), 386–399.e12. 10.1016/j.cell.2018.03.027

[15] Corces, M. R., Granja, J. M., Shams, S., Louie, B. H., Seoane, J. A., Zhou, W., Silva, T. C., Groeneveld, C., Wong, C. K., Cho, S. W., Satpathy, A. T., Mumbach, M. R., Hoadley, K. A., Robertson, A. G., Sheffield, N. C., Felau, I., Castro, M. A. A., Berman, B. P., Staudt, L. M., Zenklusen, J. C., … Chang, H. Y. (2018). The chromatin accessibility landscape of primary human cancers. Science (New York, N.Y.), 362(6413), eaav1898. 10.1126/science.aav1898

[16] Bailey, T. L., Boden, M., Buske, F. A., Frith, M., Grant, C. E., Clementi, L., Ren, J., Li, W. W., & Noble, W. S. (2009). MEME SUITE: tools for motif discovery and searching. Nucleic acids research, 37(Web Server issue), W202–W208. 10.1093/nar/gkp335

[17] Wu, T., Wu, C., Zhao, X., Wang, G., Ning, W., Tao, Z., Chen, F., & Liu, X. S. (2022). Extrachromosomal DNA formation enables tumor immune escape potentially through regulating antigen presentation gene expression. Scientific reports, 12(1), 3590. 10.1038/s41598-022-07530-8

[18] Ding, H., Hu, H., Tian, F., & Liang, H. (2021). A dual immune signature of CD8+ T cells and MMP9 improves the survival of patients with hepatocellular carcinoma. Bioscience reports, 41(3), BSR20204219. 10.1042/BSR20204219

[19] Ahmed, I., Yang, S. H., Ogden, S., Zhang, W., Li, Y., OCCAMs consortium, & Sharrocks, A. D. (2023). eRNA profiling uncovers the enhancer landscape of oesophageal adenocarcinoma and reveals new deregulated pathways. eLife, 12, e80840. 10.7554/eLife.80840

[20] Morton, A. R., Dogan-Artun, N., Faber, Z. J., MacLeod, G., Bartels, C. F., Piazza, M. S., Allan, K. C., Mack, S. C., Wang, X., Gimple, R. C., Wu, Q., Rubin, B. P., Shetty, S., Angers, S., Dirks, P. B., Sallari, R. C., Lupien, M., Rich, J. N., & Scacheri, P. C. (2019). Functional Enhancers Shape Extrachromosomal Oncogene Amplifications. Cell, 179(6), 1330–1341.e13. 10.1016/j.cell.2019.10.039

[21] Hung, K. L., Luebeck, J., Dehkordi, S. R., Colón, C. I., Li, R., Wong, I. T., Coruh, C., Dharanipragada, P., Lomeli, S. H., Weiser, N. E., Moriceau, G., Zhang, X., Bailey, C., Houlahan, K. E., Yang, W., González, R. C., Swanton, C., Curtis, C., Jamal-Hanjani, M., Henssen, A. G., … Chang, H. Y. (2022). Targeted profiling of human extrachromosomal DNA by CRISPR-CATCH. Nature genetics, 54(11), 1746–1754. 10.1038/s41588-022-01190-0

[22] Wu, S., Turner, K. M., Nguyen, N., Raviram, R., Erb, M., Santini, J., Luebeck, J., Rajkumar, U., Diao, Y., Li, B., Zhang, W., Jameson, N., Corces, M. R., Granja, J. M., Chen, X., Coruh, C., Abnousi, A., Houston, J., Ye, Z., Hu, R., … Mischel, P. S. (2019). Circular ecDNA promotes accessible chromatin and high oncogene expression. Nature, 575(7784), 699–703. 10.1038/s41586-019-1763-5

[23] Helmsauer, K., Valieva, M. E., Ali, S., Chamorro González, R., Schöpflin, R., Röefzaad, C., Bei, Y., Dorado Garcia, H., Rodriguez-Fos, E., Puiggròs, M., Kasack, K., Haase, K., Keskeny, C., Chen, C. Y., Kuschel, L. P., Euskirchen, P., Heinrich, V., Robson, M. I., Rosswog, C., Toedling, J., … Koche, R. P. (2020). Enhancer hijacking determines extrachromosomal circular MYCN amplicon architecture in neuroblastoma. Nature communications, 11(1), 5823. 10.1038/s41467-020-19452-y

[24] Widner, D. B., Liu, C., Zhao, Q., Sharp, S., Eber, M. R., Park, S. H., Files, D. C., & Shiozawa, Y. (2021). Activated mast cells in skeletal muscle can be a potential mediator for cancer-associated cachexia. Journal of cachexia, sarcopenia and muscle, 12(4), 1079–1097. 10.1002/jcsm.12714

